# Convergence of direction, location and theta in the rat anteroventral thalamic nucleus

**DOI:** 10.1101/2023.01.11.523585

**Authors:** Eleonora Lomi, Kate J. Jeffery, Anna S. Mitchell

## Abstract

Retrosplenial cortex (RSC) is a cortical region that computes heading direction from landmark information, but how it does this remains unknown. Recently we found that its two major subregions receive differential projections from two anteroventral thalamic (AV) subfields; dorsomedial (AVDM) and ventrolateral (AVVL). To probe the respective contributions of these inputs we recorded single neurons and local field potentials from AV in rats during foraging. We observed and characterized neurons modulated by theta oscillations, heading direction, and a conjunction of these. Unexpectedly, we also discovered place cells (neurons modulated by location). Consistent with the notion that AV contains two parallel subcircuits, there was a prevalence of non-conjunctive cells in AVDM, and of conjunctive and place neurons in AVVL. This integration of spatial and movement signals in AV is consistent with a thalamic role in multimodal integration and may be important for supplying the spatial information that modulates RSC directional responding.

## Introduction

We are investigating the neural circuitry underlying the mammalian sense of direction, which provides the foundation for spatial representation and navigation, as well as episodic memory functions (Aggleton et al., 2010, 1996; Harding et al., 2000; Tsivilis et al., 2008; van Strien et al., 2009; Warburton et al., 2000; Winter et al., 2015). The directional representation is supported by head direction (HD) cells, which are active when the animal faces a cell’s preferred firing direction (PFD; (Taube et al., 1990), and are located within a hierarchical system of cortical and subcortical structures comprising the extended hippocampal system (Aggleton and Brown, 1999). To understand how HD cells construct their signals it is necessary to determine what information is carried by their inputs. Here, we focused on the thalamo-cortical pathway and explored what information is carried by the thalamic inputs to the cortical HD neurons. This study was motivated by two recent findings suggesting that there are two functionally distinct (but overlapping) thalamocortical pathways. The first finding is that HD neurons in retrosplenial cortex (RSC) form two subpopulations, one in the dysgranular region (dRSC) that is more influenced by local landmarks, and one distributed across both dRSC and granular (gRSC) subregions, comprising the “classic” HD cell population that computes global heading direction of the animal (Jacob et al., 2017). The second is that the gRSC is more strongly coupled to the hippocampal theta network with a higher degree of theta-modulated firing (Lomi et al., 2021). The question arises as to what are the inputs that account for the differential behaviours of these two subpopulations of RSC neurons. To address this, we have been exploring the dense projections that arrive in dRSC and gRSC from anterior thalamus (Lomi et al., 2021; Shibata, 1993a, 1993b, 1992; Van Groen and Wyss, 1995).

Past work has focused on the anterodorsal thalamic nucleus (AD) as the main subcortical source of HD information to cortex (Goodridge and Taube, 1997; Taube, 1995; Winter et al., 2015; Yoder et al., 2015). While the majority of neurons in this dorsal region are HD cells, the picture more ventrally, in the anteroventral thalamic nucleus (AV), is slightly different: here the general consensus has been that AV is less directionally modulated and more strongly modulated by theta frequency oscillations (Albo et al., 2003; Kocsis et al., 2001; Tsanov et al., 2011, 2010; Vertes et al., 2001; Welday et al., 2011). However, evidence has also implicated the AV in directional processing, as HD cells have been reported here (Tsanov et al., 2011). The majority of these are theta-modulated – something that is not seen elsewhere in the anterior thalamus. Thus, most AV cells discharge rhythmically, and coherently with local and hippocampal theta oscillations (Albo et al., 2003; Kocsis et al., 2001; Tsanov et al., 2011; Vertes et al., 2001), and in the most dorsomedial subfield of AV (AVDM), at the border with AD, 69% of recorded units were found to express a significant directionality (Tsanov et al., 2011). Interestingly, 39% of these units were conjunctively theta- and HD-modulated, discharging rhythmic spike trains at theta frequency when the animal was facing the cell’s PFD, a property not observed along the main HD circuit via the AD (Aggleton et al., 2010; Taube, 1995). This suggests that AD and AV can be functionally differentiated.

There also seems to be functional differentiation *within* AV. Recent work, including ours, has shown that the AV-RSC projections comprise two parallel pathways (Lomi et al., 2021; Shibata, 1993a; Shibata and Yoshiko, 2015; T. van Groen and Wyss, 1990a; Van Groen and Wyss, 2003; van Groen and Wyss, 1992). One arises from AVDM and projects to gRSC, which only contains “classic” HD cells that encode the global sense of direction (Jacob et al., 2017; Lozano et al., 2017) and displays the theta hippocampal signature. This region receives projections from the hippocampus via the subiculum (Shibata, 1989; T. van Groen and Wyss, 1990a, 1990b; van Groen and Wyss, 1992; Van Groen and Wyss, 2003). These observations suggest that gRSC is more closely linked to the hippocampal spatial system. The other projection arises from AVVL and more strongly projects to the dysgranular subregion (dRSC), which is more closely linked to the visual cortex, lacks theta, and contains, as well as classic HD cells, the subpopulation of HD cells described earlier that are more driven by visual landmarks (Jacob et al., 2017) and egocentric sensory signals (Alexander et al., 2020; Alexander and Nitz, 2015). This subcircuit thus seems more sensory in nature.

The notion of parallel subcircuits is further reinforced by examination of the inputs to AV (Fig. 1). The main inputs are from medial nuclei of the mammillary bodies (MMN; Seki and Zyo, 1984; Shibata, 1992), in turn receiving inputs from the medial septum and ventral tegmental nucleus of Gudden (VTN; Shibata, 1989), both being implicated in generating hippocampal theta oscillations (King et al., 1998; Vertes et al., 2004; Vertes and Kocsis, 1997). In agreement with the idea of parallel AV pathways, MMN projections to the two AV subfields also arise from anatomically segregated populations of cells (Fig. 1 diagram; Seki and Zyo, 1984; Shibata, 1989; Shibata, 1992). Both MMN and AV receive inputs from the dorsal subiculum, which may provide a direct source of hippocampal information (Seki and Zyo, 1984; Shibata, 1994, 1993a, 1993b; van Groen and Wyss, 1992; van Strien et al., 2009).

**Figure 1:**
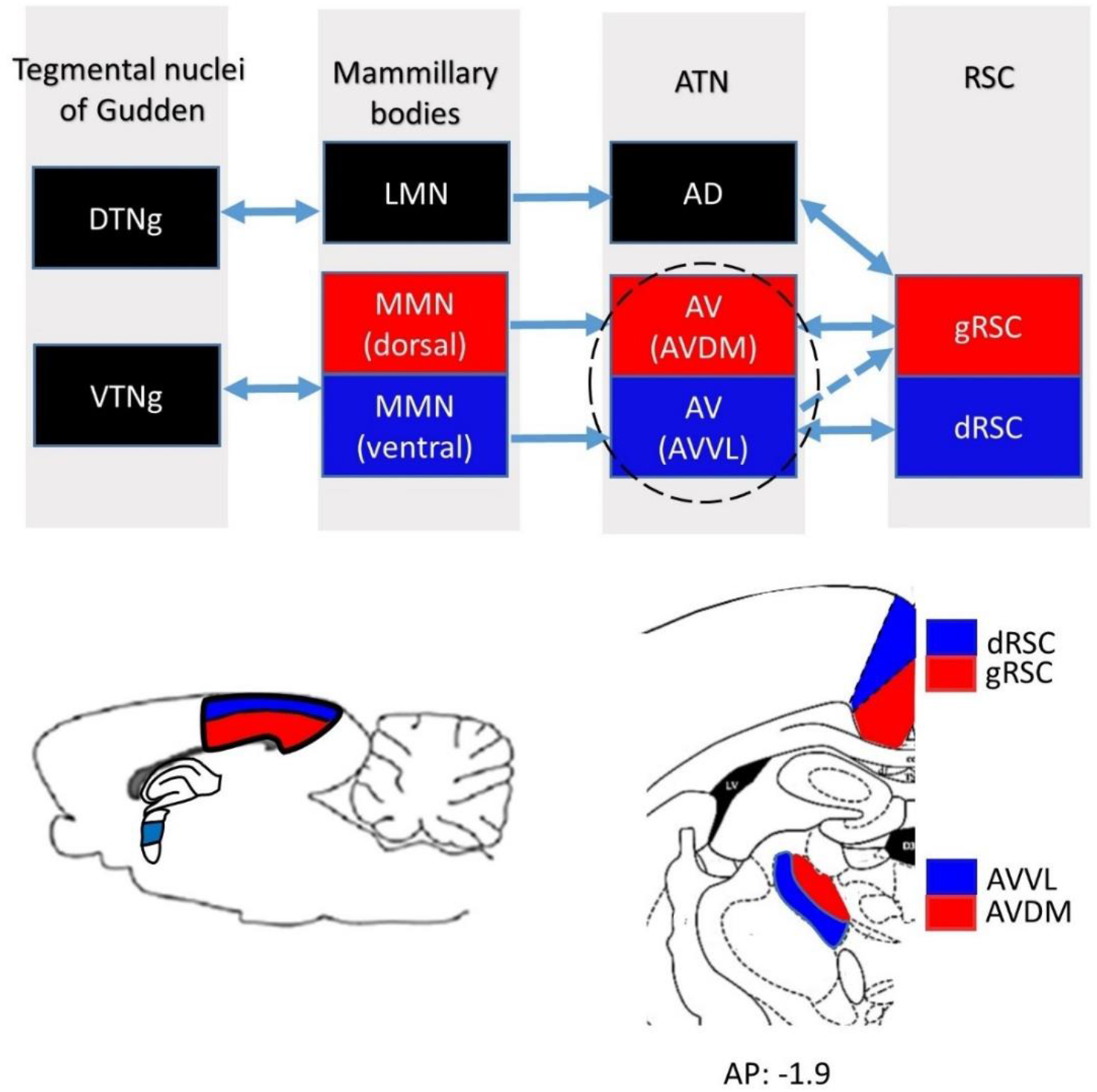
The two-pathway model of thalamocortical interactions in the head direction system. Mammillary bodies comprise a lateral (LMN) and a medial (MMN) mammillary nucleus, while the tegmental nuclei of Gudden comprise a dorsal (DTNg) and a ventral (VTNg) tegmental nucleus. Within the anterior thalamic nuclei (ATN), anterodorsal (AD) and anteroventral (AV) nuclei form separate dorsal (red) and ventral (blue) pathways that project to granular and dysgranular retrosplenial cortex (gRSC; dRSC). AV (encircled) is the site of our recordings. Bottom left, Anatomical location of RSC subregions and the three ATN: AV, AD and anteromedial (AM). AV is highlighted in blue. Bottom right, Coronal Atlas section showing RSC and AV regions, adapted from Paxinos and Watson (2007). AP, anterior-posterior level of the section.

Given this apparent two-pathway architecture, we investigated what information AV neurons convey to RSC that might help this cortical structure construct its dual landmark-dependent/sensory (via dRSC) and landmark-independent/mnemonic (via gRSC) signals. Using tetrodes, we recorded single neuron spiking and local field potentials (LFP) within the two AV subfields, AVDM and AVVL in awake, freely foraging rats. We found that both subfields contain functionally distinct subpopulations of neurons that are modulated by several spatial signals including HD and self-motion (including theta and head movement speed). When we compared between AVDM and AVVL, we did not find any difference in overall directionality nor theta modulation. However, there were fewer conjunctive cells in AVDM and correspondingly more that were only directional- or only theta-modulated. The most surprising finding was that a significant percentage of cells were place cells, showing spatially localized firing. These were found throughout both subnuclei and showed some differences from classic hippocampal place cells, but also many similarities including phase precession. Overall, our findings extend observations that anterior thalamus has both anatomically and functionally differentiated subdivisions that interact differentially with the hippocampal system.

## Results

### Multiple functional cell types in AV

For the analysis, we used the six rats for which histology indicated that electrodes sampled the AV (see Table S1). A total of 733 cells were collected from the six animals moving freely in the open field box (n = 324 in AVDM, n = 409 in AVVL; see Fig. 2 and Tables S1-S2 for breakdown of cell types and AV subfields). Of these, 504 (69%) cells were significantly theta-modulated, 280 (38%) were directionally modulated and 61 (8%) were place cells. All place cells were theta-modulated while, among the directional neurons, 158 (22%; 56.43% or directional units) were conjunctive for theta and head direction modulation. The remaining 107 cells did not fit into any clear classification.

**Figure 2:**
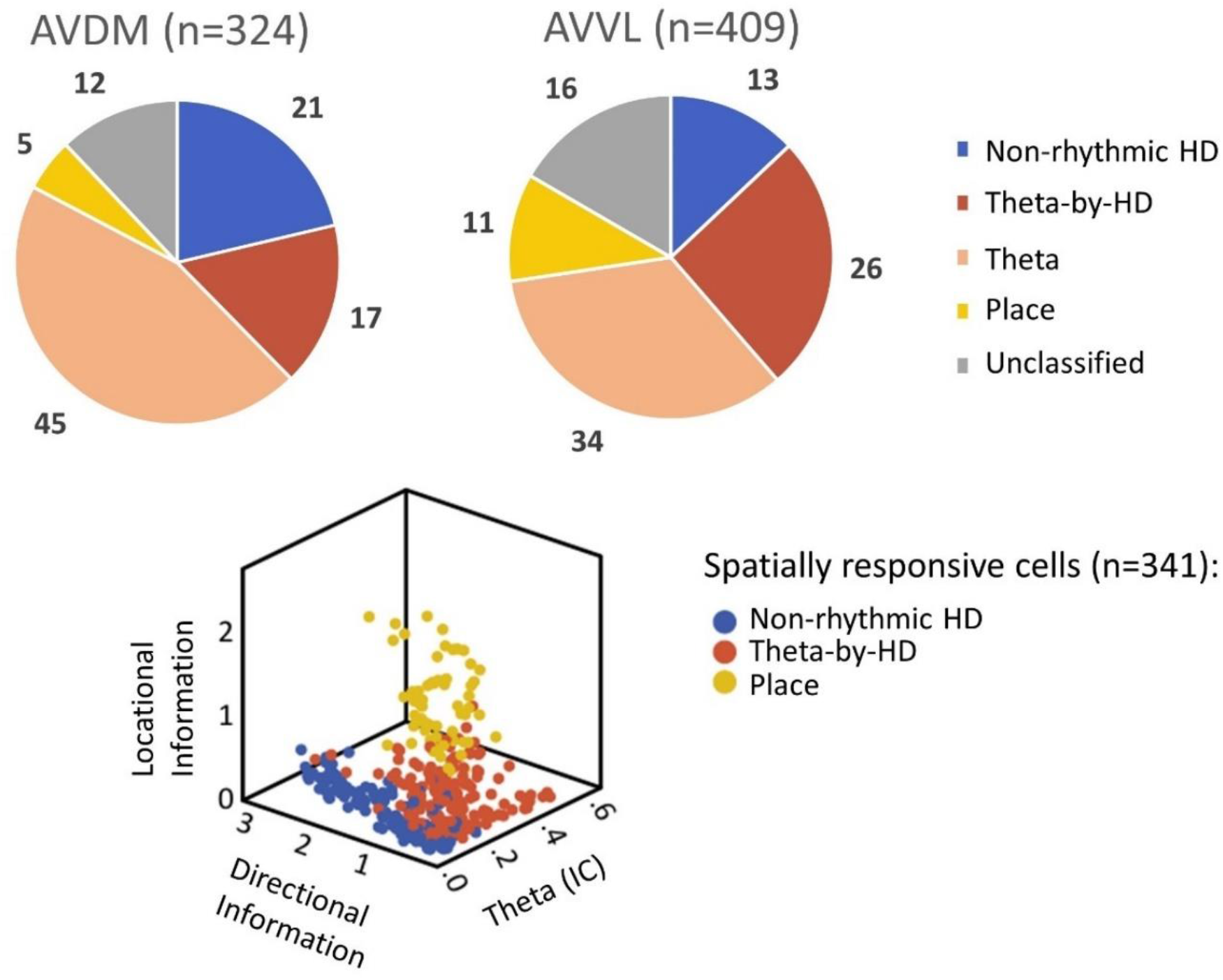
Different anatomical distributions of functional cell types within two anteroventral thalamic (AV) subfields. Top, proportion of cells in the two AV subfields: dorsomedial (AVDM) and ventrolateral (AVVL). Cell numbers are as follows (AVDM vs AVVL): non-rhythmic HD (122), 69 vs 53; Theta-by-HD, 53 vs 105; place, 17 vs 44; Theta, 146 vs 139; Unidentified, 39 vs 68. ** = 0.01 significance, Chi-squared test. There was no difference in the anatomical location of the unclassified units with no spatial/temporal correlates. Bottom, the three groups of spatial/directional cells were well segregated in a scatterplot along three independent parameters.

When we compared between AVDM and AVVL subfields we did not find any difference in overall directionality (38% and 39%, respectively) nor in theta modulation (61% and 60%, respectively). However, there were fewer conjunctive cells in AVDM (16% vs. 26%; χ2 = 9.28, *p* = 0.0023) and correspondingly more that were only directional (21% vs. 13%; χ2 = 9.06, *p* = 0.0029) or only theta-modulated (45% vs. 34%; χ2 = 9.33, *p* = 0.0022). These cell types (Fig.2) and their properties are examined in turn, below.

We start with the spatial/directional cells (examples in Fig. S1-S4). As expected, based on previous work, HD cells (n = 280; 38%) were found throughout the AV subfields: AVDM = 122; 44%; AVVL = 158; 65%. These consisted of two subtypes, with distinct directional properties (Fig. 3A; analysis reported below). The most surprising observation was that a subset of cells from 5/6 animals (n = 61; 8.32%) were place cells (Fig. 7A), which have not been reported previously in anteroventral thalamus. These cells were recorded throughout both AV subfields (AVDM = 17; 5%; AVVL = 44; 11%) and formed single (Fig. S3) or multiple (Fig. S4) discrete firing fields. Fields were always highly compact and stable across trials (Fig. 8B. In two animals, we recorded seven place cells simultaneously on the same tetrode as one or two Theta-by-HD cells, across six different days (examples in Fig. 8C), confirming that recorded place cells were part of the local AV circuit and not ectopic cells dragged down from the overlying hippocampus (which anyway would have been unlikely, given the distance).

**Figure 3:**
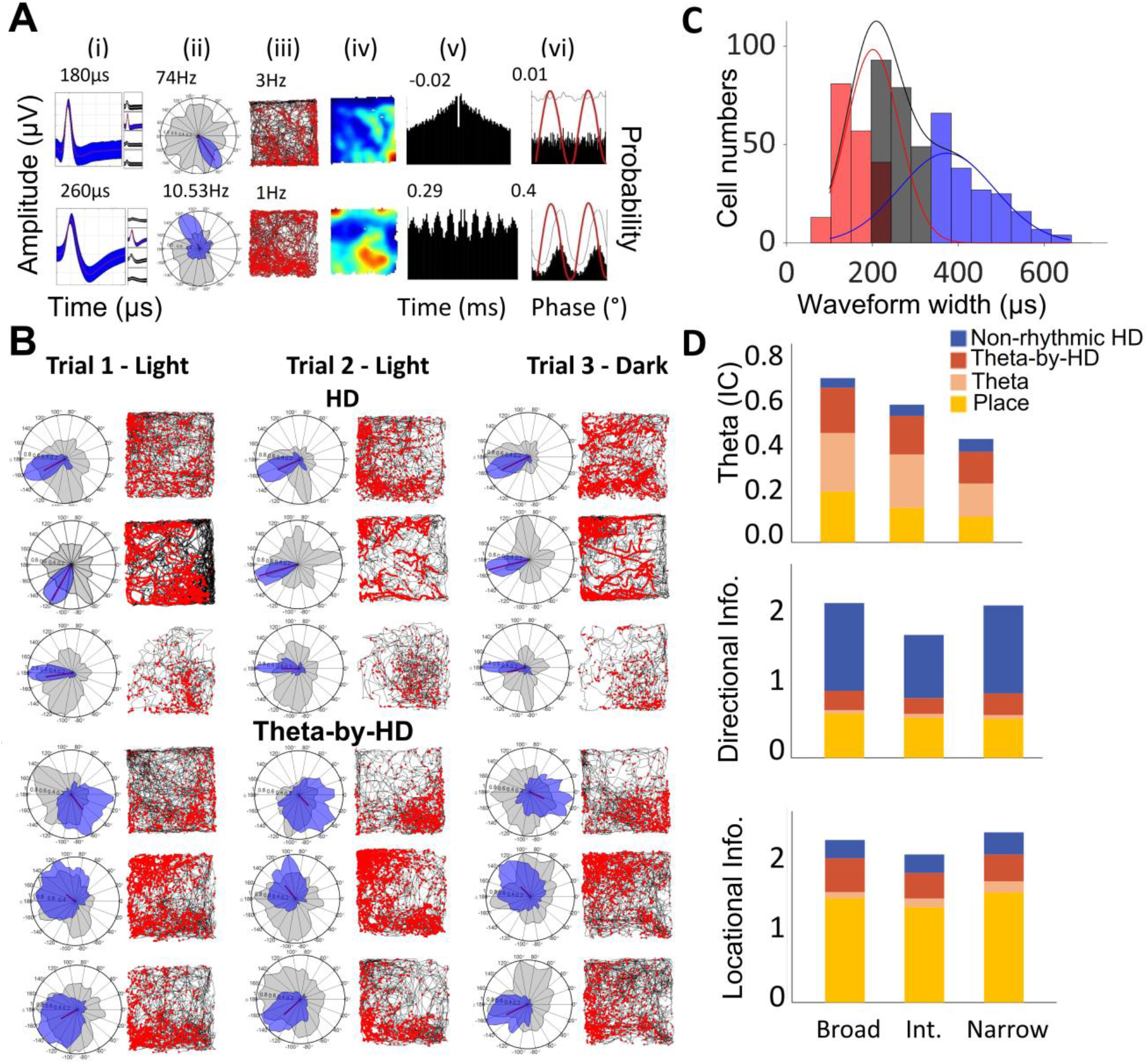
Two main types of directional cells in the anteroventral thalamus (AV): theta-rhythmic and non-rhythmic. (A) An example of each is presented. For each cell the following plots are shown: (i) Average tetrode spike waveform. Red line is the mean, shaded area the standard deviation. Average peak-to-trough width (μs) are reported on top. (ii) Polar plot of firing rate (Hz) vs head direction (degrees) in blue and dwelltime (seconds) vs head direction (HD; degrees) in grey. Peak firing rates are reported. (iii) Spikemap and (iv) ratemap with mean firing rate (Hz), showing uniform distribution of spikes across the box. (iv) Temporal autocorrelogram implemented between ±500ms with index of rhythmicity (IR). (v) Double-plotted histogram of spike frequency relative to theta phases. The thin line is the corresponding smoothed density estimates (see Methods). The red line is the sine wave representing two theta cycles. x-axis, degrees between 0-360; y-axis, probability. (B) Examples of HD and Theta-by-HD recorded across three consecutive trials. Spikemaps and polar plots of firing rate vs 360° of HD (blue) and dwelltime vs HD (grey) are shown for each trial (each column is a trial; third is in darkness). (C) Frequency distribution of peak-to-trough waveform width for AV cells. x-axis, μs; y-axis, frequency count. Gaussian fits for the different modes (waveform types) are displayed: broad (blue), narrow (red), and intermediate (grey). See also Fig. S5. (D) For each cell type, distribution on theta, directional and locational properties are presented as stacked bar charts, with cell types as grouping variable. Theta is more likely to occur in broad compared to narrow and intermediate waveforms. In contrast, directionality is more likely to occur in narrow and intermediate waveforms compared to broad. Locational selectivity is equally represented.

Having provided an overview of the functional cell types recorded, we next took each of these cell types in turn and investigated their physiological properties and firing correlates (summarised in Table S3).

### Waveform analysis reveals different neuronal subtypes

We asked whether cell groups identified in AV presented differences in their waveform shape, as this can be indicative of the AV cell typology (Henze et al., 2000; McCormick et al., 1985). Approximately half of the spatial units (193/341, 56.6%) had a narrow peak-to-trough waveform (<250μs): non-rhythmic HD, 58.20%; Theta-by-HD, 53.80%; place cells, 56.60% (Table S3). Average width did not differ between these groups (KW test, H(3) = 49.54; *p* < 0.001; all *p* > 0.5) but all groups had a narrower width than the theta cells (all *p* < 0.01), for which only 32.28% of spikes were narrow.

We plotted the distribution of waveform widths (Fig. 3C) and fitted a two-Gaussian model. Values fell into three distinct clusters, narrow (n = 192; 30.70%), broad (n = 183; 37.08%), and intermediate (n = 221; 32.22%). Cells were discarded if spike waveforms did not reach time for repolarization (the global minimum of the curve occurs before the peak, n = 30; 5.03%). The calibrated Hartigan’s Dip Test discarded the null hypothesis of unimodality (DIP = 0.04; *p* < 0.001). We compared the firing characteristics of cells in the three groups. The odds of finding a cell with a significant theta component in the broad spiking group was 5.8x (CI: 1.95, 6.56) and 3.21x (CI: 1.76, 5.85) higher, respectively, than in the narrow and intermediate groups, respectively (Fisher exact tests; all *p* < 0.0001). In contrast, cells with a significant HD component were 3.39x (CI: 2.201, 5.24) and 3.22x (CI: 2.11, 4.92) more likely to be found in the narrow and intermediate groups, respectively, than in the broad one (all *p* < 0.001). The odds of finding a cell with a place component was not significantly different than one for all the three groups (all *p* > 0.05; Fig. 3D). The variability of spike width (Fig. S3-S4) argues against place cells being axon spikes.

Peak-to-trough waveform width was greater for AVDM (330.24 ± 9.34μs) than AVVL neurons (284.83 ± 7.25μs; two sample WRS, z = 4.26; *p* < 0.0001), as expected from their morphology, and may be a contributing factor for the variable waveform shapes reported above.

### Place cells show evidence of complex spike bursting

We next looked at the basic temporal patterns of firing (Fig. S5A-B). Place cells had the lowest mean rate, followed by Theta-by-HD cells and then non-rhythmic HD and theta cells, whose rate did not differ from each other (KW test; H(3) = 78.96; *p* < 0.001; HD vs Theta-by-HD, *p* = 0.02; HD vs Theta, *p* =1; HD vs place, *p* < 0.001; place vs Theta-by-HD, *p* < 0.01; place vs Theta, *p* < 0.01; Theta-by-HD vs Theta, *p* < 0.01; Table S3). As expected from the waveform analysis above, there was a weak, but significant, negative correlation between peak-to-trough width and mean rate (Pearson’s r = -0.1; *p* = 0.01).

AV cells presented differences in a number of other temporal firing properties. Place cells contained the highest proportion of bursting units (69%), while Theta-by-HD (47%) and theta (33%) cells contained similar proportions of bursting and non-bursting units and non-rhythmic HD cells only contained a marginal proportion of bursting units (7%). Proportions differed significantly from each other (Chi-square tests, all *p* < 0.001). This is visible as a high central peak for the autocorrelations of place cells (Fig. S3-S4), suggestive of complex spike bursting. Place and Theta-by-HD cells showed theta modulation, burst firing and a shorter ISI compared to non-rhythmic HD cells, which only fired single spikes that were mostly unrelated to theta (Table S3). Fig. S5C shows examples of bursting and non-bursting units, and the distribution of peak ISI for the identified spatial cells.

Consistent with the larger concentration of place cells in AVVL (reported below), spike bursting propensity was higher in AVVL (peak ISI: 16.15 ± 1.28ms; burst index: 0.09 ± 0.00) compared to AVDM (peak ISI: 22.33 ±2.00; two sample WRS, z = - 3.49; *p* < 0.001; burst index: 0.08 ± 0.01; z = -2.87; *p* = 0.0041). There was no difference in mean firing rate between AVDM (3.62 ± 0.29Hz) and AVVL (3.32 ± 0.19Hz = -2.874, *p* = 0.68).

### Oscillatory entrainment of unit activity by the local LFP theta

We examined oscillatory activity in the local LFP, as a prelude to looking at theta modulation of spiking and found notable activity in the Type-1 theta frequency range (6-12Hz; average power: 3.49 ± 2.53; average theta-delta ratio: 1.32 ± 1.65; average frequency: 8.79 ± 0.64Hz). Across all six animals, we found a strong modulation by the animals’ running speed (Fig. 4; r = 0.90 ±0.13 for theta power and 0.87 ± 0.18 for theta frequency).

**Figure 4:**
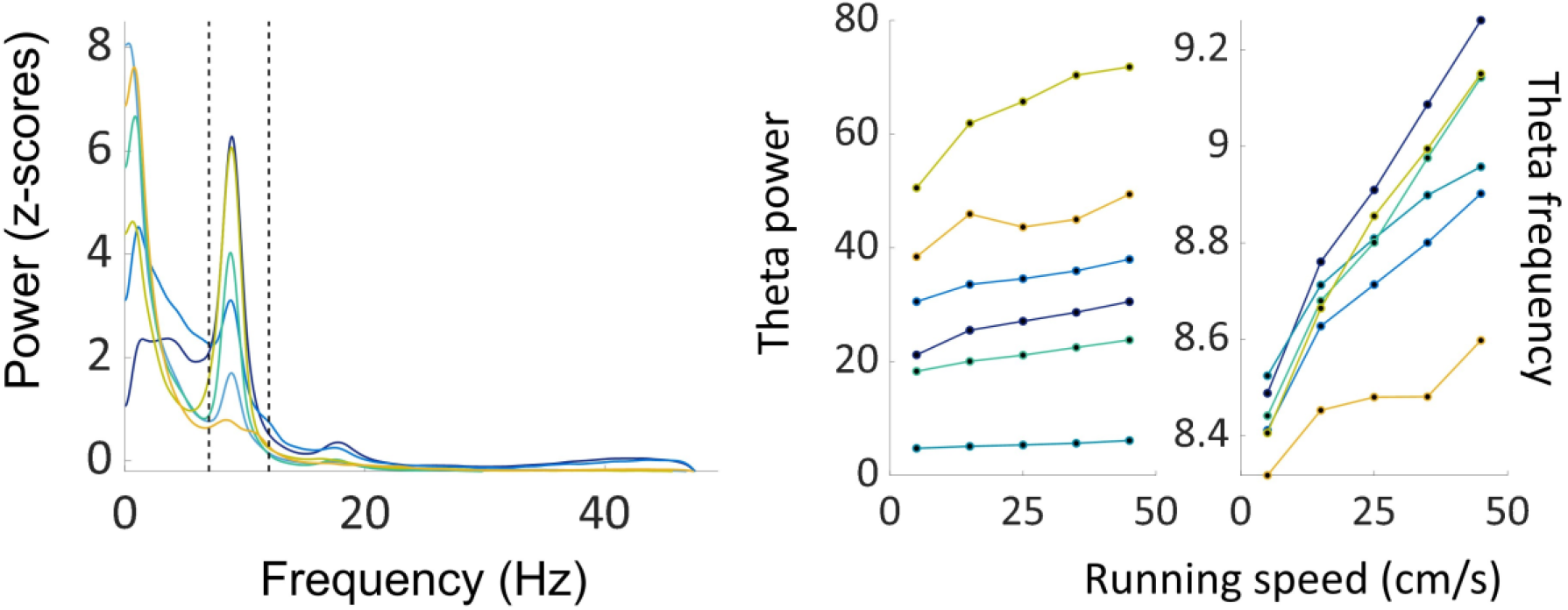
Theta oscillations locally recorded in anteroventral thalamus (AV) are modulated by running speed. Left, average normalized power spectrum across all trials, color coded by animal (n=6). Dotted lines define theta range (6-12Hz). Right, average speed-power and speed-frequency relationships across all trials for each animal, color coded as in A.

We next analyzed the theta-band properties of single cells. Theta rhythmicity and coupling were assessed by computing an index of rhythmicity (IR) and coupling (IC), respectively (Fig. S6), and then we looked at two other known theta-associated phenomena: theta phase precession and theta skipping.

Theta-by-HD cells, place cells and theta non-directional cells were theta phase-locked, with the same average IC and IR, all being significantly higher than non-theta rhythmic HD cells (IC; KW test; H(3) = 206.54; IR; H(3) = 180.86; all *p* < 0.001; Table S3). We also looked at a phenomenon known as theta skipping, in which the autocorrelogram reveals a second theta peak larger than the first, due to a subpopulation of cells that fired on alternate theta cycles (Brandon et al., 2013). We saw theta skipping in 29.11%, n = 46/158) of Theta-by-HD cells (Fig. S7).

For cells that fired locked to the local LFP theta (which is phase-coupled and precedes hippocampal theta by 7-10ms; Tsanov et al., 2011), we computed the preferred theta phase and plotted these values in circular histograms (Fig. 5A). For all cells, spikes were aligned to descending theta phases. The theta peak was defined as 0° and place cells fired closer to the peak (76.02°; R-vector = 0.59; Rayleigh test, z = 21.65, *p* < 0.0001) while Theta-by-HD cells (124.02°; R-vector = 0.67; z = 69.82; *p* < 0.0001) and theta non-directional cells (115.42°; R-vector = 0.50; z = 71.58) fired slightly after place cells, closer to theta trough. Both differed significantly from place cells (two-samples WW tests, Theta-by-HD vs Theta, *F* = 1.96; *p* = 0.16; Theta-by-HD vs place, *F* = 31.48; Theta vs place, *F* = 18.15; all *p* < 0.0001), suggesting sequential, phase-offset activation between these cells. The earlier average phase preference for place cells might be due to the steeper precession of firing phases relative to theta (shown below).

**Figure 5:**
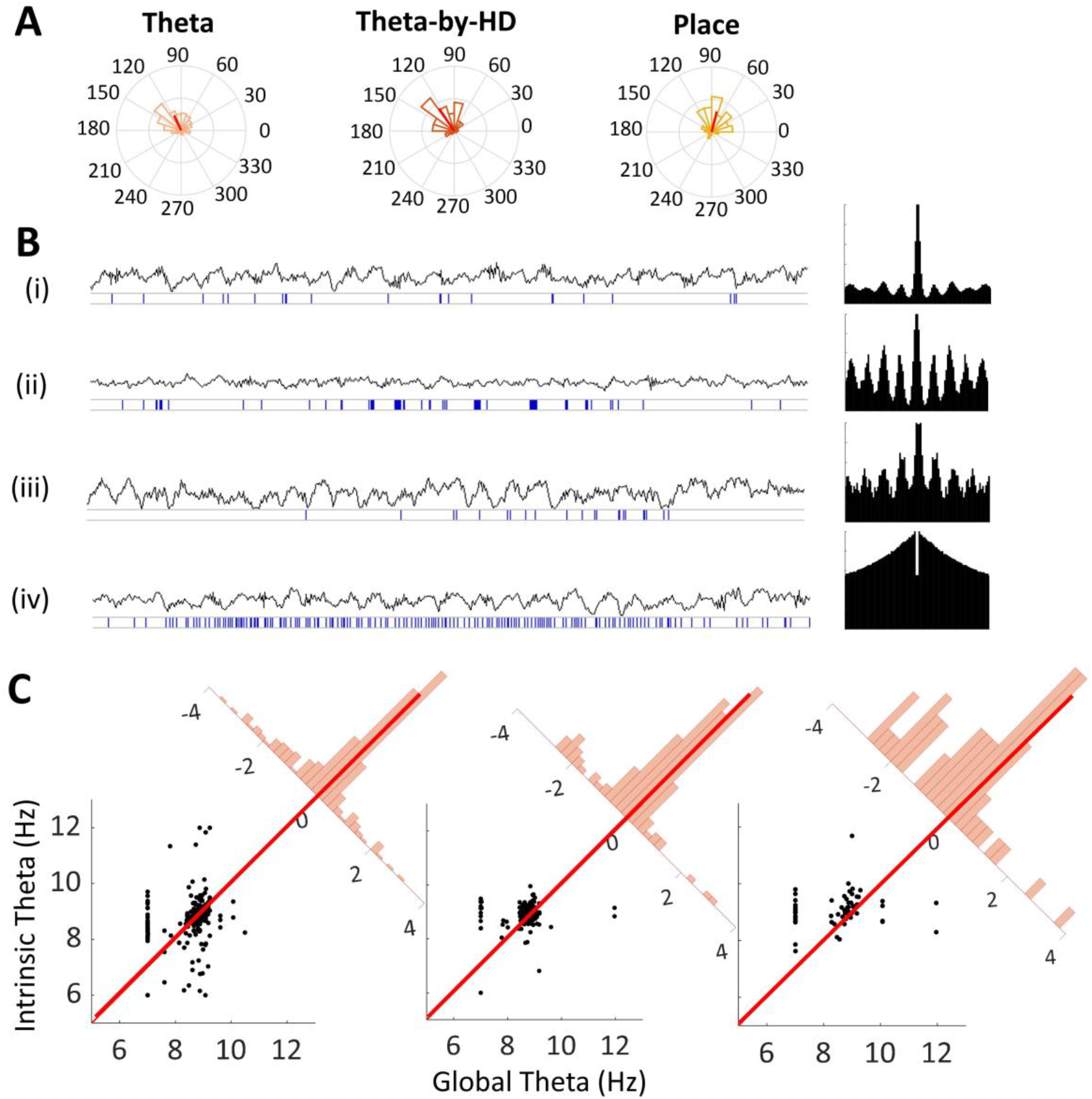
Periodic firing of AV units during theta states. (A) Circular histograms of preferred theta phases for theta-modulated cells: theta non-directional (n = 285), Theta-by-HD (n = 158), place (n = 61). Red line is the R-vector pointing toward the mean preferred theta phase. Radius = 0.3. All cells fire between 0-180° (descending theta phases) but place cells fire slightly earlier, just after the peak (closer to 0). No difference between firing phases of theta and Theta-by-HD cells. (B) Examples of unit recordings and raw LFP trace recorded from the same electrode (for 3s of recordings). Cells are in the following order: (i) Theta non-directional; (ii) Theta-by-HD; (iii) Place; (iv) HD non-theta cells. The corresponding temporal autocorrelogram for each cell, between ± 500ms, is shown to the right. Theta cells with no spatial properties mostly fired single spikes (not part of a train) while Theta-by-HD and place cells discharged rhythmically grouped spike trains (groups of 2-3 spikes with short inter-spike intervals) at theta frequencies. Theta-modulation is not visually detectable in the traces of non-rhythmic HD cells. Place cells show theta phase precession: spikes occurred slightly earlier with successive theta cycles. Note theta-skipping for the Theta-by-HD cell. (C) Analysis of theta phase precession for the phase-locked units. Intrinsic vs global theta frequency (Hz), shown as scatterplots in the same order as in A. Each marker is a cell; if a marker is above the diagonal line, the cell was phase precessing. Corner histograms display the distance of each marker from the diagonal. x-axis, distance to diagonal (a negative distance means the cell falls above the diagonal). Place and Theta-by-HD cells: distributions are toward negative values. Theta non-directional cells: more centrally aligned (weaker phase precession).

For the theta-modulated neurons, we also looked at the phenomenon of phase precession, first reported in place cells (O’Keefe and Recce, 1993), in which successive spikes from a bursting neuron occur at progressively earlier phases of the simultaneously recorded theta LFP. This can be revealed if average intrinsic theta frequency of a neuron is slightly faster than global theta (Fig. 5B-C). Comparing intrinsic oscillation frequency to the LFP frequency allows to derive a convenient count of precessing neurons. We found evidence of phase precession in place cells (intrinsic vs global frequency: 9.09 vs 8.54Hz; two-sample t-test, *t*(60)= -3.60, SD = 1.20, *p* < 0.0001) and Theta-by-HD cells (intrinsic vs global frequency: 8.9 vs 8.63Hz; *t*(157) = -4.08, SD = 0.82, *p* < 0.0001). Theta cells did not show strong phase precession (intrinsic vs global frequency: 8.76 vs 8.64Hz; *t*(284) = -2.46, SD = 0.87, *p* = 0.01). Overall, 77.05% (47/61) place cells and 63.92% (101/158) Theta-by-HD cells phase precessed. Proportions did not differ from each other (χ^2^ = 3.46, *p* = 0.063), but were higher than theta cell proportion (Theta-by-HD vs Theta, χ^2^ = 4.96, *p* = 0.026; place vs Theta, χ^2^ = 11.89, *p* < 0.0001).

We finally compared between the two AV subfields. Previously, we found that theta activity differed between the two RSC subregions, being more present in gRSC than dRSC (Lomi et al., 2021). In contrast, there was no theta difference between AV subfields (AVDM vs AVVL; two-sample WRS tests, IC: 0.17 ± 0.01 vs 0.14 ± 0.01, z = 0.21, *p* = 0.834; TS index: -0.004 ± 0.01 vs -0.007 ± 0.01, z = 0.05, *p* = 0.960).

### Two types of directional firing

Having examined the temporal firing characteristics of AV units, we now move to their directional properties. 288 (38%) cells had significant directionality in their firing. As mentioned earlier, both theta-modulated (Theta-by-HD) and non-theta (HD) cells were found in both AV subfields, but the non-rhythmic HD cells were more prevalent in AVDM (χ2 = 9.06, *p* = 0.003) and the theta HD cells more prevalent in AVVL (χ2 = 9.28, *p* = 0.002).

There were other differences between the two cell types (Fig. 6A-B). Theta-by-HD cells displayed less-precise directional firing compared to non-rhythmic HD cells: Theta-by-HD cells showed lower R-vectors (0.62 ± 0.02 vs 0.33 ± 0.02; two-sample WRS test, z = -9.74), broader tuning widths (94.04 ± 3.0 vs 132.30 ± 1.15; z = -9.74) and smaller peak rates (27.06 ± 2.92 vs 5.20 ± 0.03Hz; z = 6.59). The concentration parameter, which gives a measure of how directionally compact the firing is (like the sparsity score used for place cells), was lower for Theta-by-HD cells (0.72 ± 0.03 vs 2.53 ± 0.19; z = 9.74; all *p* < 0.001).

**Figure 6:**
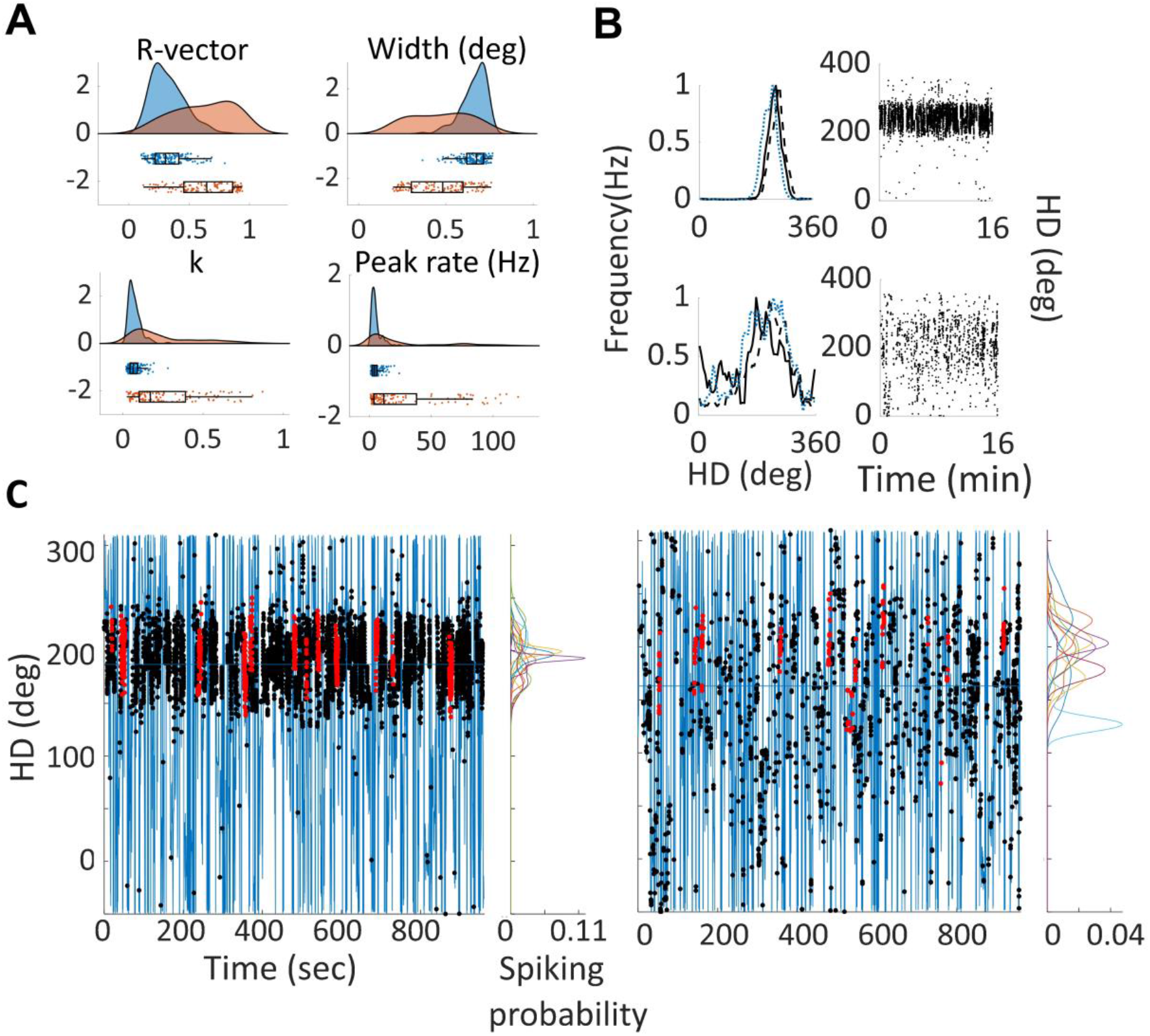
Two directional signals: a broad one operating at theta frequency, and a narrow one independent of theta. (A) Distribution of tuning curve parameters, shown as raw data + boxplot + split-violin plot. k, concentration parameter. Theta-by-HD cells show lower directional specificity that non-theta HD cells. (B) Peak normalized tuning curves for non-theta HD (top) and Theta-by-HD (bottom) cells, calculated over short timescales: solid line, first 3 minutes; dashed line, last 3 minutes; blue line, remaining trial duration. Theta-by-HD cell tuning curves are broader since the start. For each cell, the corresponding scatterplot of HD vs time are shown. (C) For the same cells in B, spatial plots of unsmoothed directional heading as a function of time are plotted (blue line), overlaid at the appropriate times with spikes (black markers). Red markers are spikes emitted during full head turn events through the cells’ preferred firing direction (when the cells fired >50% of its intra-trial peak rate plus spikes within +/-50°). Kernel density estimates (KDE) for each successive event are plotted on the right and show a larger variability for Theta-by-HD cells.

To explore why Theta-by-HD cells had broader tuning curves, we looked at the tuning across a trial and found that Theta-by-HD cells shifted directions more during a trial, which could have led to a broader tuning curve when collapsed across the trial (Fig. 6C). Specifically, Theta-by-HD cells displayed larger jitter in the HD data when they fired more than 50% of intra-trial peak rate (SEM of peak location: non-rhythmic HD, 13.66 ± 0.60°; Theta-by-HD, 16.59 ± 0.67°; two-sample T-test, t(244) = -3.16, sd = 7.21, *p* = 0.002). In contrast, KDE peaks for Theta-by-HD and non-rhythmic HD cells had an equivalent mean width, suggesting that the range of directional firing was actually the same (non-rhythmic HD, 33.9 ± 0.75; Theta-by-HD, 35.59 ± 0.68; t(252)= -1.67, sd = 8.02, *p* = 0.1). SEM of peak width was also the same (non-rhythmic HD, 4.52 ± 0.22; Theta-by-HD, 5.22 ± 0.28; t(244) = -1.92, sd = 2.87, *p* = 0.06).

However, not all of the tuning curve broadening in Theta-by-HD cells could be explained by drift because at small time intervals, when drifting is small, we saw residual broadening. This was investigated by constructing plots of firing rate vs HD for the first 3 minutes of recordings (Fig. 6B). Average tuning width continued to be broader for Theta-by-HD (133.09 ± 1.33°) compared to non-rhythmic HD cells (97.43 ± 2.80°; two-sample WRS test, z = -9.56, *p* < 0.0001). Likewise, tuning curves remained broader when the animal was turning CW, CCW or when still (see Supplementary Information; Table S5), suggesting that a larger variability of heading directions might also be a fundamental property of these cells.

Cells maintained stable PFDs across trials. Tuning curve correlations were high across both Light-Light trials (non-rhythmic HD, 0.89 ± 0.19; Theta-by-HD, 0.81 ± 0.01) and Light-Dark trials (non-rhythmic HD, 0.89 ± 0.02; Theta-by-HD, 0.79 ± 0.02), with no difference between the two (two-sample KS test; non-rhythmic HD, ks-stat = 0.07; Theta-by-HD, ks-stat = 0.1; *p* > 0.5; Fig. 7A). Observed and control distribution of correlation coefficients differed significantly from each other (Fig. 7B) in both conditions for non-rhythmic HD cells (two-sample KS test, Light-Light, ks-stat = 0.94; Light-Dark, ks-stat = 0.94; all *p* < 0.0001) and Theta-by-HD cells (Light-Light, ks-stat = 0.92; Light-Dark, ks-stat = 0.95; all *p* < 0.0001). However, in line with the greater intra-trial drift for Theta-by-HD cells (shown above), correlations were higher for non-rhythmic HD cells across both light and dark conditions (two-sample KS test; Light-Light, ks-stat = 0.58; Light-Dark, ks-stat = 0.59; *p* < 0.0001). For both cell types, absence of vision did not affect coding of orientation (bits/spike; WSR test, non-rhythmic HD, z = -0.4631; Theta-by-HD, z = -0.23; *p* > 0.1) and overall firing rate (non-rhythmic HD, z = 1.66; Theta-by-HD, z = 2.40; *p* > 0.01 (Fig. 7C).

**Figure 7:**
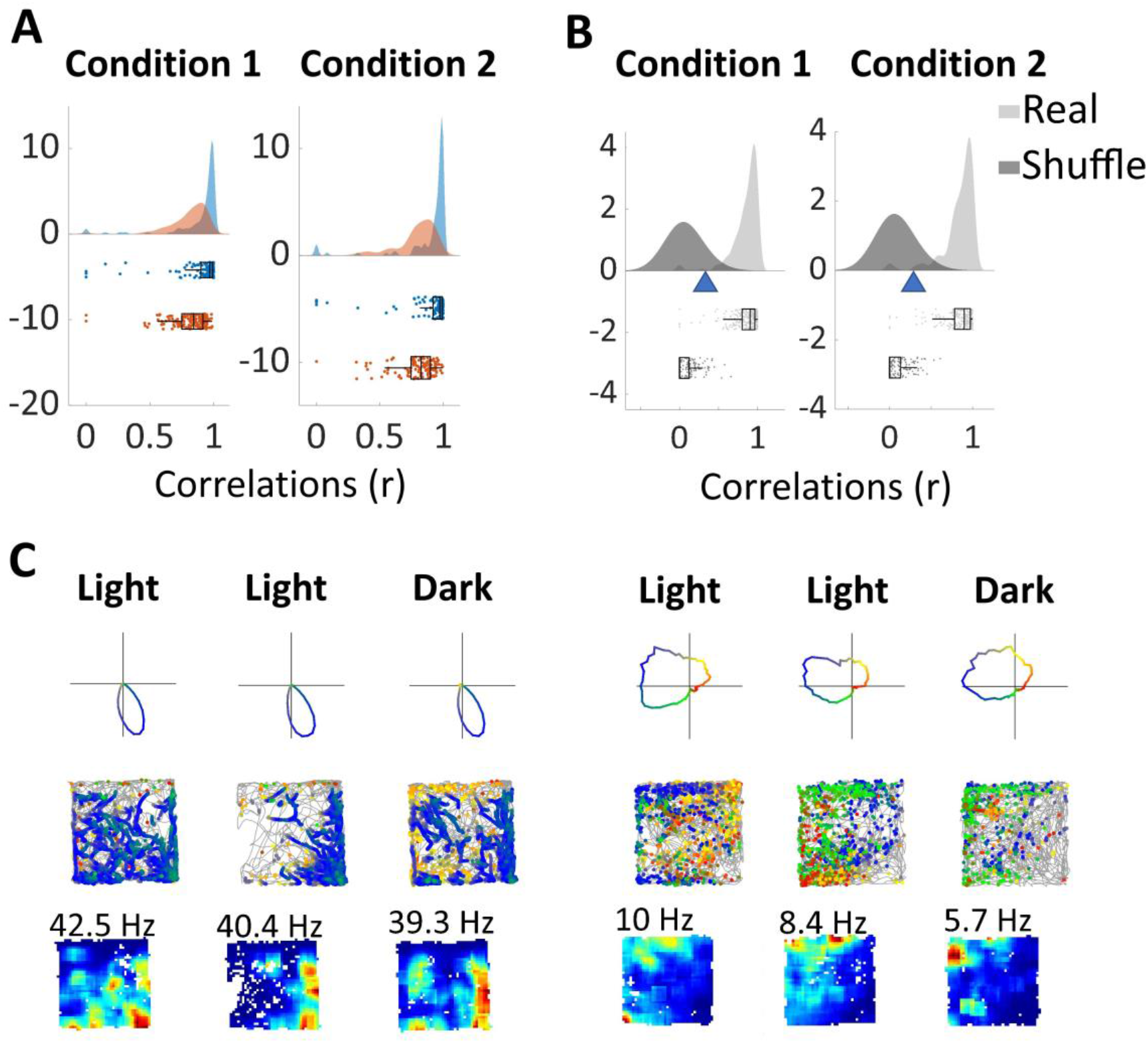
Directional firing is similar across Light-Light (Condition 1) and Light-Dark (Condition 2) trials. For each Condition: (A) Distribution of trial pair-specific tuning curve correlations for non-rhythmic HD (blue) and Theta-by-HD (red) cells. (B) Distribution of correlation coefficients for cells in A, compared to shuffle distributions, blue triangles represent the 95^th^ percentile shuffling. No difference between Conditions. (C) Examples of non-theta HD (left) and Theta-by-HD (right) cells recorded across the three trials. Each trial shows the following: Top, directionally color-coded polar plot, with direction of peak firing in blue, and the other directions following a color gradient changing every 90°; Middle, directionally color-coded spikemap, with spikes colored according to their direction overlaid on the path of the rat (highly smoothed in 20ms bins); Bottom, ratemap with peak rate reported. See Fig. S10 for place cell examples.

### Spatial firing

Cells were examined for the presence or absence of spatial firing patterns. As mentioned, surprisingly, 61 cells (8%) were found to have place fields (Fig. 8A). These AV place cells primarily formed single place fields in the 90 × 90cm box (n= 5/61 had 2-3 fields regularly spaced), like hippocampal neurons recorded in similar sized environments (Fenton et al., 2008; Harland et al., 2021). Some Theta-by-HD cells (n = 20/158; 13%) also passed the place cell criteria, although spatial tuning was much less specific than for the place cells and the cells always fired over more than 20% of the recording arena. This is reflected in the ratemaps, which occasionally displayed broad patches of spatially inhomogeneous firing (Fig. S2-S9). No non-rhythmic HD cells met the place cell criteria.

**Figure 8:**
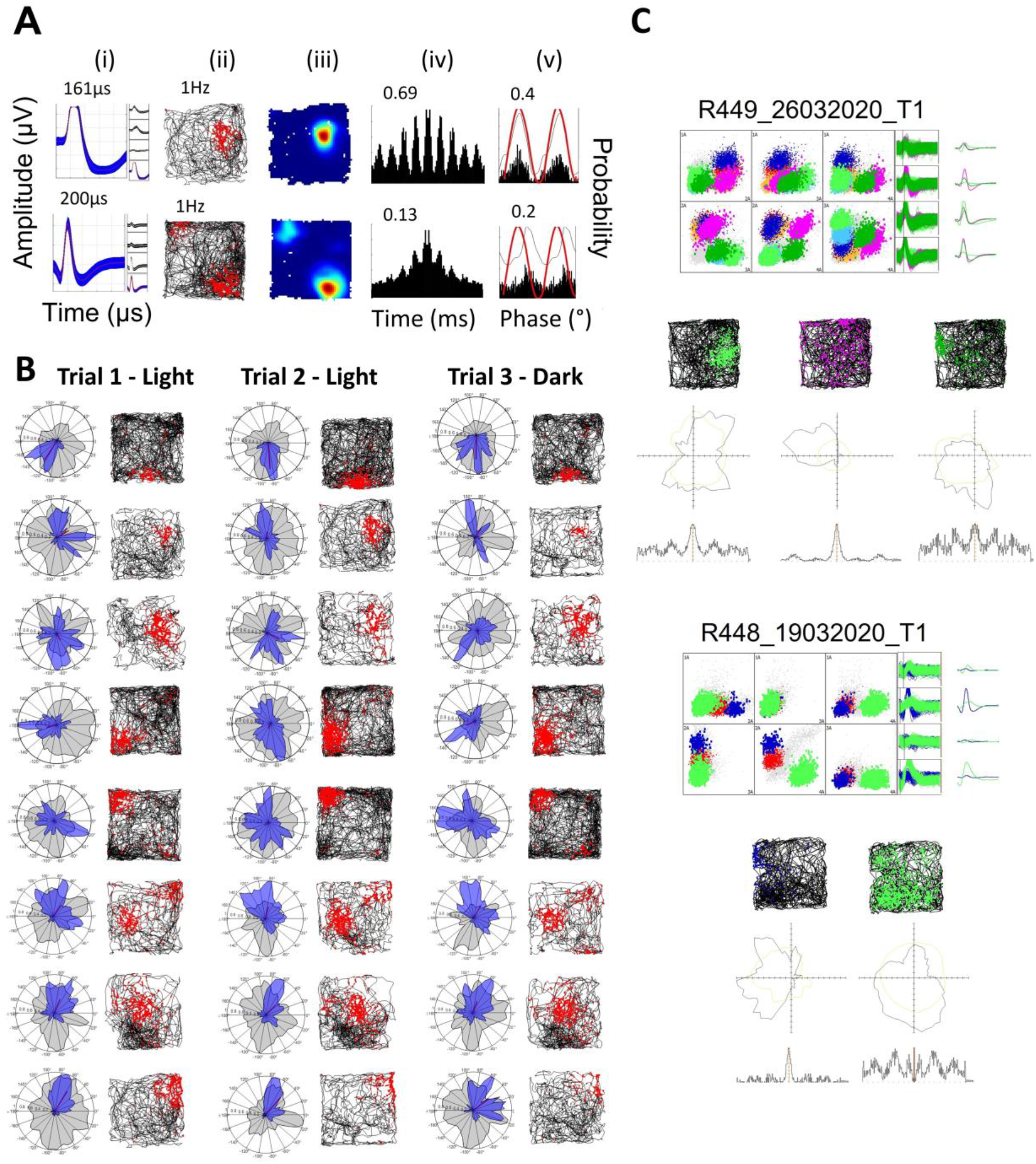
Place cells are highly stable across trials and can be co-recorded on the same tetrode with directional neurons. (A) Examples of place cells with single and multiple firing fields; plots as in Fig. 3A. (B) Stability of place fields across trials and in darkness; plots as in Fig. 3B. See also Fig. S8-S9. (C) Examples of two recording trials in which place cells were co-recorded on the same tetrode with a Theta-by-HD cell. The tetrode cluster space and the average tetrode waveform for each cluster are shown. Spikemap, polar plot of firing rate (Hz) vs HD (degrees) and temporal autocorrelogram between ± 300ms are shown for each unit.

For the place cells, we then looked at the metric properties of the firing fields. In our data, place fields were smaller and more spatially compact (mean field area: 717.67 ± 44.78 cm^2^; sparsity: 0.15 ± 0.01) compared to previous reports from dorsal hippocampal regions (fields size typically above 1000 cm^2^, Mizuseki et al., 2012; sparsity around 0.24, Zhang et al., 2014). We found no difference in place field size between AVDM (655.76 ± 76.50 cm^2^) and AVVL (741.59 ± 54.68 cm^2^; two-sample t-test, *t*(59) = -0.86, SD = 87.62, *p* = 0.39). The fields were characterized by high locational information (1.51 ± 0.06 bits/spike) and locational selectivity (15.15 ± 1.13), and low sparsity (0.15 ± 0.01), compared to HD cells (locational information: 0.35 ± 0.02 bits/spike; selectivity: 4.24 ± 0.14; sparsity: 0.52 ± 0.9; Table S3). Post-hoc tests showed that Theta-by-HD cells were intermediate in locational information content between non-rhythmic HD and place cells (Table S3).

We looked at whether there were AV subfield differences in the cells’ spatial properties. Consistent with the larger concentration of place cells in AVVL, shown above, locational modulation was higher in AVVL (locational information: 0.26 ± 0.02 bits/spike) compared to AVDM (0.26 ± 0.02 bits/spike; two sample WRS, z = -3.32, *p* < 0.001). Directional modulation was the same in both (AVDM, directional information: 0.33 ± 0.03 bits/spike; R-vector: 0.25 ± 0.01; AVVL, directional information: 0.32 ± 0.02 bits/spike; z = -0.45, *p* = 0.653; R-vector: 0.26 ± 0.01; z = -1.45, *p* = 0.146).

While testing in the dark did not cause place remapping (Fig. 8B-S10), nor a change in global mean firing rate (z = -0.96, *p* > 0.1), locational information was reduced in Dark compared to Light conditions (paired sample WSR test, z = 4.16, *p* < 0.0001). It is possible that a small place field drift occurred in the absence of visual anchoring.

### Velocity sensitivity and anticipatory firing

Finally, we analyzed movement correlates, in linear and angular domains. Most of the cells showed positive responses to speed (n = 194/269; 72.12%) and AHV (n = 32/44; 72.73%), that is firing rates increased as the animal ran faster, and its head turned faster in both turning directions (see example of a negatively tuned place cell in Fig. 9). We first look at linear, then angular speed correlates. Most cells (269/341; 78.89%) met the criterion for linear speed tuning. All cell types were different from zero (one-sample WSR test; non-rhythmic HD, z = 9.59; Theta-by-HD, z = 10.90; place cells, z = 6.79, all *p* < 0.0001). Equal proportions of speed-tuned units were observed in each cell group (Chi-squared tests; all *p* > 0.1; Fig. 9; Table S3) and tuning strength did not differ between groups (average absolute s-scores: non-rhythmic HD, 0.56 ± 0.02; Theta-by-HD, 0.52 ± 0.02; place cell, 0.55 ± 0.03; KW test; H(2) = 1.771, *p* = 0.41).

**Figure 9:**
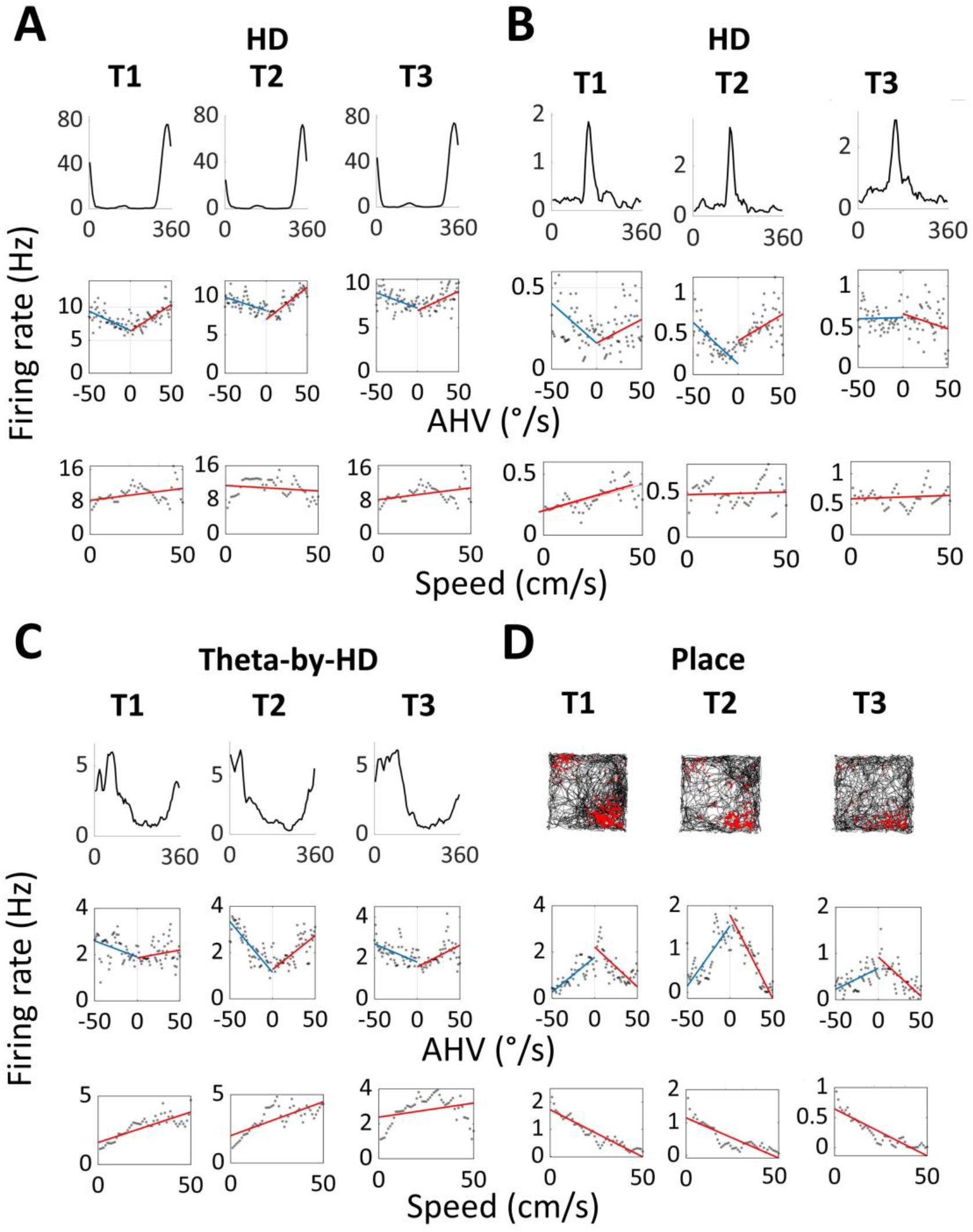
Spatial cells are modulated by linear and angular head movements. Examples of (A-B) non-rhythmic head direction (HD), (C) Theta-by-HD and (D) place cells showing conjunctive modulation by running speed and angular head velocity (AHV). Each cell was recorded across three consecutive trials. Trial type is at the top (T1, T2, T3; T2 was in darkness). For each cell, the following plots are shown, from left to right: tuning curve of firing rate (Hz) vs HD (degrees) or spikemap for the place cell; scatterplot of instantaneous firing rate (Hz) vs AHV in 2° /s bins; scatterplot of instantaneous firing rate vs running speed in 2cm/s bins. Linear regression lines were fitted to derive the slope and the y-intercept of the rate-AHV (clockwise (CW), red; counter-clockwise (CCW), blue) and the rate-running speed relationships. All conjunctive cells displayed the same direction of linear and angular activation (both positive or both negative).

In the angular domain, likewise, equal proportions of AHV-tuned units were observed in each group (44/341; 12.90%; Chi-squared tests; all *p* > 0.1; Table S3) and strength of AHV activation did not differ across cell types (average absolute AHV-scores: non-rhythmic HD, 0.22 ± 0.01; Theta-by-HD, 0.24 ± 0.01; place cell, 0.25 ± 0.02; KW test; H(2) = 2.31, *p* = 0.315) and differed slightly but significantly from zero (one-sample WSR test; non-rhythmic HD, z = 9.58; Theta-by-HD, z = 10.90; place cell, z = 6.791, all *p* < 0.0001), suggesting weak, yet significant, AHV activation. For almost all of the AHV-tuned cells (42/44; 95.45%), firing rate modulation was symmetric (no effect of turning direction; paired-sample WSR test, z = 2.06, *p* = 0.04). Most angular-tuned neurons were conjunctively tuned to linear speed (n = 39/44; 88.64%).

We then compared across AV subfields: proportions of cells responding to speed or AHV did not differ between AVDM and AVVL (see values Table S3; all *p* > 0.05).

We next looked at anticipatory firing. HD cells can fire in relation to future, rather than present, HD by integrating information about current HD with the velocity at which the head is turning (Blair et al., 1997). Non-rhythmic HD cells were more anticipatory (anticipatory time interval (ATI) = 41.7 ± 2.56ms) than Theta-by-HD cells, the firing of which was only slightly lagging the animal’s HD (−17.3 ± 6.64ms; Fig. 10A-B). ATI values differed from each other (two-sample WRS test, z = 7.05, *p* < 0.0001). In relation to this, the CCW-CW difference angle was positive for non-rhythmic HD (9.71 ± 1.32°) and negative for Theta-by-HD cells (−9.54 ± 1.54; Kuiper two-sample test, k = 9312, *p* = 0.001). Non-rhythmic HD cell, but not Theta-by-HD cell values, differed significantly from zero (one-sample WSR test; HD, z = 8.85, *p* < 0.0001; Theta-by-HD, z = -1.82, *p* = 0.069).

**Figure 10:**
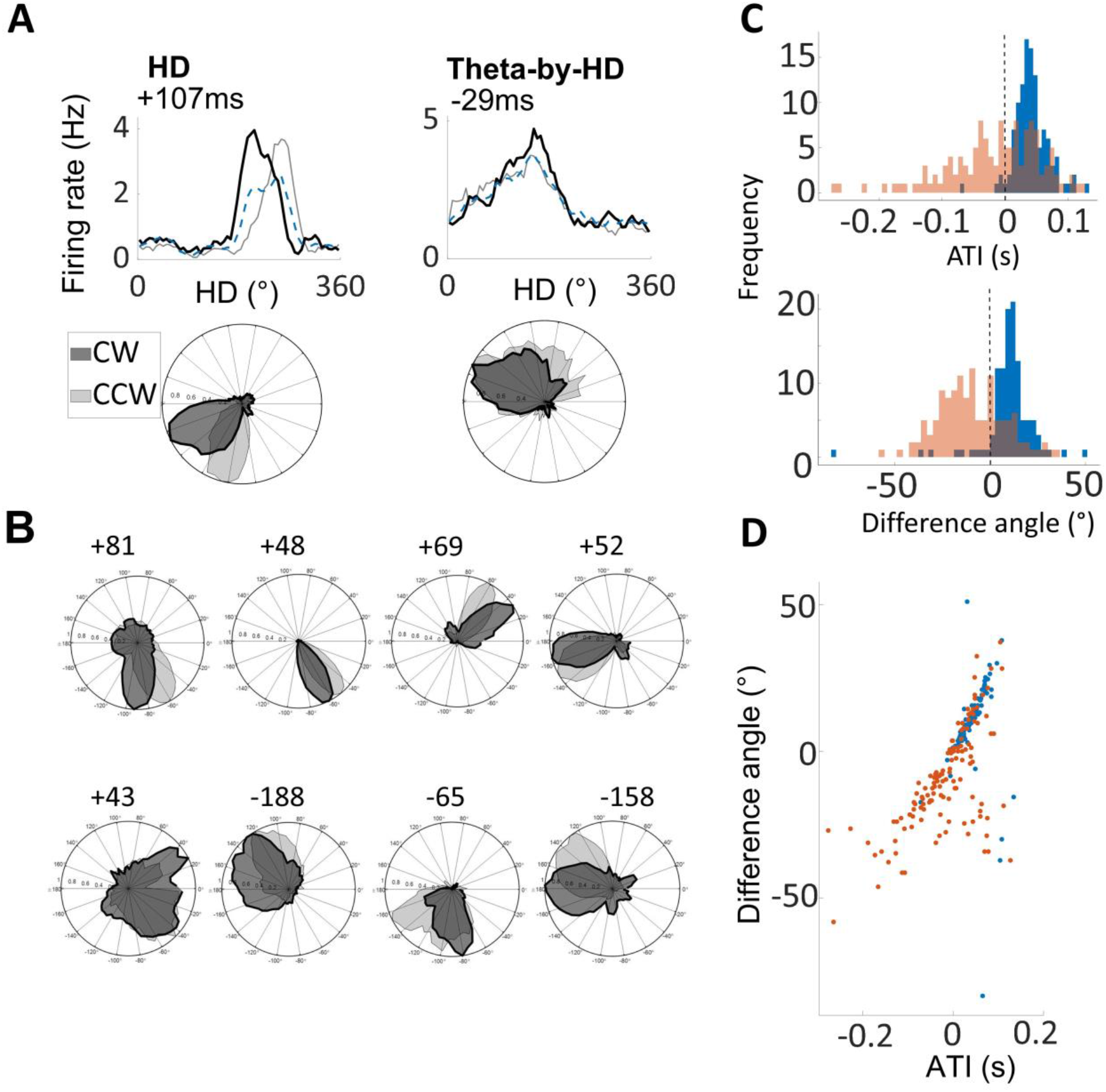
Non-rhythmic head direction (HD) cells are more anticipatory than Theta-by-HD cells. Tuning curves decomposition analysis used to derive a cell’s anticipatory time interval (ATI). (A) Tuning curve decomposition analysis for a classic HD (non-rhythmic) and a Theta-by-HD cell. For each cell, the tuning curve of firing rate (Hz) vs head direction (HD; degrees) is shown as smoothed histogram (top) and as polar plot normalized to 1Hz (bottom). Tuning curves were decomposed into clockwise (CW; black) and counter-clockwise (CCW; grey). The standard tuning function (blue) includes all spikes, including those emitted during immobility. ATI for each cell is reported. (B) More examples of the cells in A. Theta-by-HD cells are less anticipatory (smaller CW-CCW angular separation) and maintain their broader tuning curves also when decomposed into CW and CCW (see analysis in Supplementary Information and Table S5). Note that negative ATIs mean tuning functions shift in opposite directions during CW and CCW turns compared to positive ones. (C) Frequency distribution of ATIs and CCW-CW difference angles, compared between classic HD (blue) and Theta-by-HD cells (red). For both measures, Theta-by-HD show a greater variability while classic HD cells are skewed toward positive values. (D) Positive correlation between these measures. Markers represent cells, color coded as in C. The more a cell is anticipatory, the more the tuning curve shifts in opposite directions during CW and CCW turns.

Theta-by-HD cells displayed a greater amount of variability (Fig. 10C), with some cells lagging and others anticipating the actual direction: ATIs were positive for 96% of non-rhythmic HD (107/112) and for only 56% of Theta-by-HD cells (59/128). Nevertheless, correlation between implant location and average ATI values did not reach significance (Pearson’s *r* = -0.01, *p* = 0.87) and there was no difference in the proportion of Theta-by-HD cells with negative ATIs between the two subfields: AVDM and AVVL (Chi-square test for differences in proportion; χ^2^ = 0.24, *p* = 0.625). The depth of the cell within AV may also be a contributing factor. As expected, there was a positive correlation between ATI and difference angle measures (Pearson’s r = 0.63, *p* < 0.0001; Fig. 10D).

We additionally computed ATI values using the optimal time-shift method, which relies on the standard tuning function alone (see Methods) and obtained values that closely matched those estimated with the above cross-correlation method. Mean optimal shifts for non-rhythmic HD (width, 40.33 ± 5.70ms; peak rate, 35.40 ± 3.40ms) were higher than Theta-by-HD (width, -10.51 ± 6.42ms; peak rate, -7.6 ±7.10ms; two-sample WRS tests; peak rate, z = 4.62; width, z = 5.75, all *p* < 0.0001). Non-rhythmic HD cells were more anticipatory (87% positive for peak rate, 79% for width), while Theta-by-HD cells exhibited greater variability (40% for peak rate, 43% for width).

### Histology

Cell locations were estimated based on the reconstruction of the electrode tracks from histology and the record of screw turns that lowered the electrodes each day, starting from a known depth. Fig. 11 illustrates examples of brain sections in which the electrode track is visible. The electrodes tip is marked and represents the deepest recording site. Two rats (R449, R222) had AVVL-only recordings; one (R762) had AVDM-only recordings. The remaining three (R448, R651, R652) had both AVDM and AVVL recordings. See details on recording sites in Table S2. To be sure that the place cells were indeed located in AV, in two animals we induced high-frequency current lesions at the end of recordings (R448, R449) through electrodes that recorded both a Theta-by-HD and a place cell on that day. The lesion was verified to be located in AV; no damage to hippocampal and parahippocampal tissue was visible in the histological slides that contained these tracks, and there was no evidence of any pulling-down of hippocampal tissue into the anterior thalamic region.

**Figure 11:**
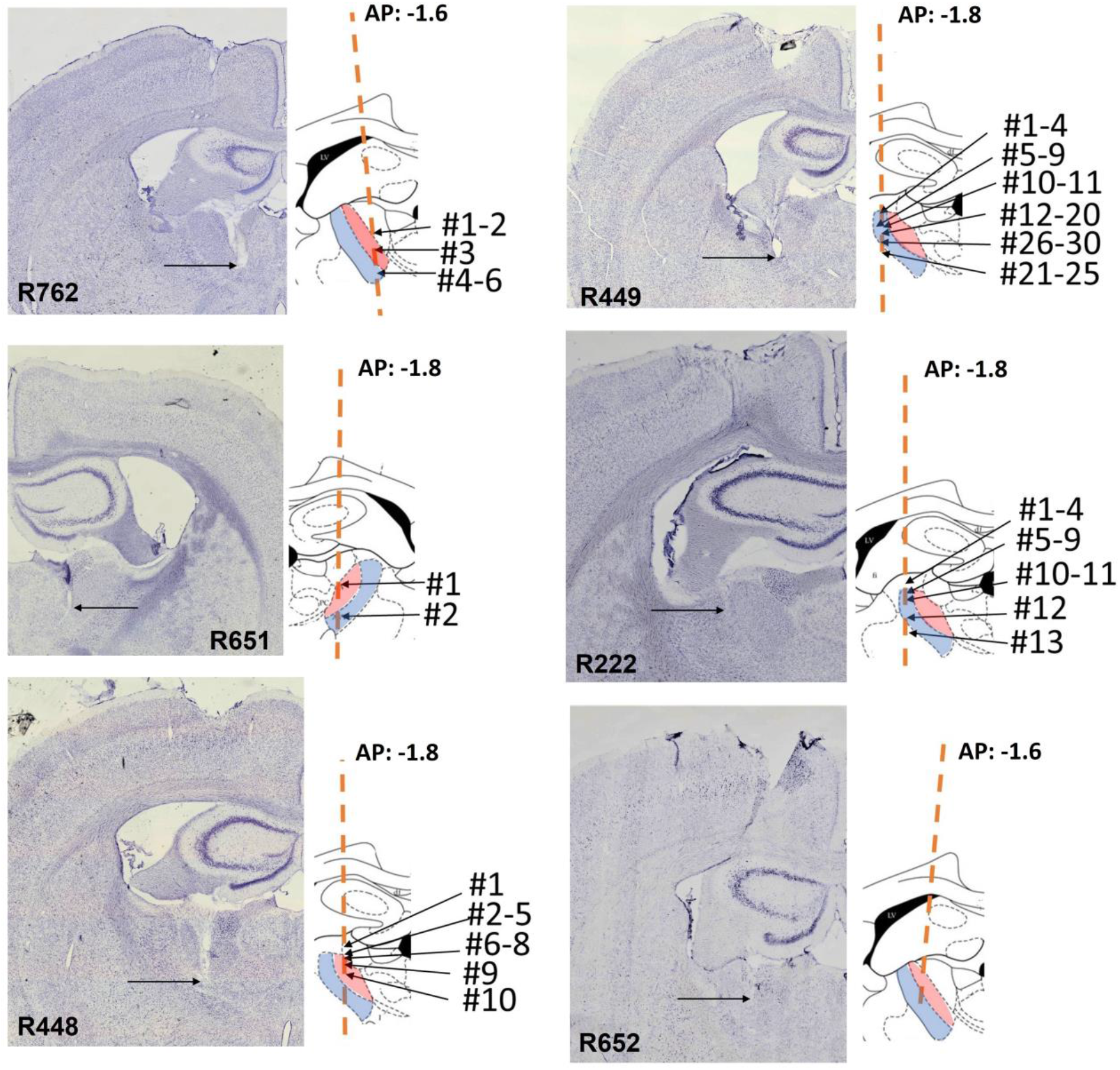
Histology of anteroventral thalamus (AV) single unit recordings. Photomicrographs of the Nissl-stained coronal sections through the recording site, for the six implanted rats. Insets on the right show schematic line drawing of AV subfields (AVDM, red; AVVL, blue) for each animal, adapted from Paxinos and Watson, 2007. Location of place cells are marked for five rats for which place cells could be recorded. Place cell location is reconstructed on the atlas section and is indicated by numbers. Orange line is the estimated track direction. Anterior-posterior (AP) location of each atlas section is indicated (−1.6 or -1.8mm from Bregma). Abbreviations: AVDM, dorsomedial anteroventral thalamus; AVVL, ventrolateral anteroventral thalamus.

## Discussion

We investigated the single-neuron processing of spatial, directional, temporal and movement parameters in anteroventral thalamus (AV), motivated by recent findings that the AV’s two subfields, dorsomedial (AVDM) and ventrolateral (AVVL), project differentially to two subregions of RSC that have distinct directional and theta signalling properties (Lomi et al., 2021). We replicated the previous AV findings of Tsanov et al. (2011) of two types of signal: a theta signal, reflected in the burst firing of neurons at the ∼8 Hz theta frequency, and a directional signal reflected in the head-direction-specific firing. These two signals interacted to produce three types of neuronal activity: theta-modulated but non-directional, directional but non-theta-modulated, and a mixed subtype (in the Theta-by-HD cells) expressing both. Our first novel finding is that, unexpectedly, we also found a population of place cells, with properties similar to those of hippocampal place cells but with different firing field metrics. The second is that AV cells, including place cells, came in two flavors, broad and narrow, although these did not conform to the spiking classes (constituting another difference from hippocampal recordings; (Csicsvari et al., 1999; Mizuseki et al., 2012; Mizuseki and Buzsáki, 2013; Wilent and Nitz, 2007). And finally, when we compared these properties between AVDM (projecting preferentially to gRSC) and AVVL (projecting preferentially to dRSC) we found several differences; notably, that the AVVL contained a higher proportion of theta-modulated cells.

We discuss these findings and their implications in more detail below, relating them to the notion of two distinct pathways from AV to RSC.

### Cell-type heterogeneity

We begin with our observations of physiological heterogeneity of neurons in the AV, manifest as differences in waveform shape, temporal firing patterns, directional firing patterns and spatial firing. In terms of waveform shape, we found two distinct clusters, reflecting a narrow spike and broad spike respectively, with the broad type being more prevalent in AVDM. This is consistent with their respective magnocellular (large cells) and parvocellular (small cells) morphology, in which the magnocellular cells have been found to be more prevalent dorsomedially and the parvocellular ones ventrolaterally (Gabriel, 1993; Shibata and Yoshiko, 2015).

We did not, however, find functional correlates of these differences. In terms of temporal firing properties, place cells and Theta-by-HD cells fired rhythmic spike trains at theta frequencies, with or without bursting, while non-rhythmic HD cells only fired single spikes unrelated from theta. For the spatially modulated neurons, namely Theta-by-HD, place and non-rhythmic HD cells, we did not find any waveform correlates, with around half of neurons in each functional type being broad and the other half narrow. Place modulation was equally represented in these two categories. All theta-modulated cells preferentially fired on descending theta phases. Place and Theta-by-HD cells showed firing phase precession in time, which was stepper than for the non-directional theta units. These functional cell types were differently distributed between AVVL and AVDM. Altogether, then, it seems that there are a range of physiologically distinct cell types in this region, but they do not conform clearly to functional categories.

### Directional tuning

We then examined the sub-class of neurons that showed above-chance directional tuning. These fell into two distinct categories: theta-modulated with broad tuning, and non-theta-modulated with narrow tuning. The latter resembled cells typically seen along the main HD circuit (AD; (Taube et al., 1990), including in their tendency to anticipate the animal’s head direction by a small amount. By contrast, Theta-by-HD cells showed reduced directional tuning and occasionally (n=20) showed some conjunctive spatial tuning, which was much weaker than in place cells, and did not show anticipatory firing. Results indicate the presence in AV of two separate HD networks: a global continuous HD network attractor, possibly inherited from corticothalamic projections, and a theta-modulated HD network that may receive more indirect, or weaker, inputs from the main HD system.

### Spatial tuning

Perhaps the most unexpected finding from this experiment was the observation of a very clear spatial signal, manifest as the occurrence of “place cells” – that is, cells firing in a restricted region of the environment. These cells resembled the classic hippocampal place cells in a number of respects, and indeed when we first saw them, we thought they might *be* hippocampal cells, since the dorsal-most part of the hippocampus overlies the anterior thalamus. However, examination of the histological slides did not reveal any evidence of explantation of hippocampal tissue into the AV; also spatial firing was observed throughout the dorsoventral extent of the electrode’s passage through the AV thalamus (and indeed was more pronounced ventrally), and in most cases, co-occurred with recordings from Theta-by-HD cells, which do not occur in hippocampus (Leutgeb et al., 2000; Royer et al., 2010).

Furthermore, the spatial firing fields were smaller than have been reported for hippocampal place cells in a similar-sized environment (Casali et al., 2019; Mizuseki et al., 2012; Mizuseki and Buzsáki, 2013). And finally, the credibility of this observation is supported by the fact that place cells have been reported in other thalamic subnuclei: notably the nearby anteromedial thalamic nucleus (AM, Jankowski et al., 2015), but also the nucleus reuniens (Jankowski et al., 2014) and paratenial nucleus (Jankowski et al., 2015). It is unlikely that our electrodes penetrated AM: they were more lateral, and the co-occurrence of Theta-by-HD cells was not reported in AM. However, the place cells were similar in other respects, such as the compact nature of the place fields and narrow waveforms. The findings of place cells in a variety of thalamic nuclei, as well as in the anterior claustrum (Jankowski and O’Mara, 2015), invites a more detailed investigation of whether the thalamus is a source or recipient of hippocampal place information, of how the signal is mixed with other signals such as head direction and theta, and of what this processing might be for.

### How do AV cells acquire their specific response properties and what are they for?

For the purposes of this discussion, we speculate on answers to the above questions as they pertain to the AV subfield. The first question is how the neurons in AV generate their functional response characteristics. The AV receives inputs from three main spatial systems: the HD system, via thalamo-cortico-thalamic projections; the hippocampus, via dorsal subiculum projections; and the brainstem theta system, via the ascending MMN pathway. In turn, the main outputs are to RSC and hippocampal formation (Shibata, 1994, 1993a, 1993b, 1992; Van Groen and Wyss, 1995; Witter et al., 1990). To explain the functional cell types recorded, we now suggest a neural network in which AV contains four main cell subpopulations. Some AV cells receive a strong descending input from the main HD network in RSC (from the subcortical HD ring attractor; Winter and Taube, 2014) and thus are non-rhythmic HD cells; another subpopulation receives ascending MMN inputs and thus are theta-modulated cells lacking spatial properties; a third subpopulation receives inputs from both the HD signal from non-rhythmic HD cells (signalling orientation) and the theta input (signalling movements) and thus become Theta-by-HD cells. A fourth receives spatial inputs, possibly via the hippocampal-subicular pathway (Christiansen et al., 2016).

The time delay between Theta-by-HD and non-rhythmic HD cells (approximately 50 ms) suggests that Theta-by-HD cells might arise from the output of many non-rhythmic HD cells with similar PFDs. This convergence could account for above-chance directionality together with lower directional specificity than is seen in single HD cells. Spatial inputs from place cells may also provide a weak input to the Theta-by-HD cell signal, given that few Theta-by-HD cells yielded weak locational specificity.

How can AV inherit spatial selectivity? AV has strong connections with limbic regions that signal spatial location, including the dorsal subiculum (Christiansen et al., 2016; Jankowski et al., 2013; Perry and Mitchell, 2019; Seki and Zyo, 1984; Shibata, 1993a; van Strien et al., 2009), where place cells exhibit broad place fields (Frost et al., 2021; Lever et al., 2009; Poulter et al., 2021; Sharp, 1999; Sharp and Green, 1994). Projections from the subiculum to ATN are bilateral to AVVL and AM, while AVDM receives only ipsilateral projections (Seki and Zyo, 1984). This is consistent with our result of fewer place cells recorded from AVDM compared to AVVL. However, it is interesting to note that ATN is required to establish place cell firing in subiculum, but not in CA1 (Calton et al., 2003; Frost et al., 2021). Given our findings, this effect could be mediated by direct AV place cell discharge to the subiculum (Shibata, 1993a; Shibata and Yoshiko, 2015; Thomas van Groen and Wyss, 1990, 1990; van Strien et al., 2009; Wright et al., 2010).

### The two AV-RSC pathways are functionally distinct

We now come back to one of the primary motivations for this investigation, which is to see whether the differential directional properties occurring in dysgranular vs. granular RSC might be accounted for by differences in their anterior thalamic afferents. Based on our recent anatomical study (Lomi et al., 2021), we compared the properties of neurons recorded in AVDM vs. AVVL. We found no difference in either theta modulation or directional modulation. However, what differed was the conjunction between these signals, since AVVL had more conjunctive neurons. We found the following differences (summarized in Fig. 2 and Table S3): (1) Classic HD (non-rhythmic) cells were more prevalent in AVDM (21% vs. 13%); (2) Theta non-directional cells were more prevalent in AVDM (45% vs. 34%); (3) Theta-by-HD cells were more prevalent in AVVL (26% vs. 16%); (4) Place cells were more prevalent in AVVL (11% vs. 5%).

These differences suggest that the AV-RSC pathway is not homogenous but may comprise two distinct subcircuits, differing in the extent to which theta and spatial neural signals are integrated. One, via AVDM-gRSC, is predominantly involved in the exchange of theta and HD information; the other, via AVVL-dRSC, is more involved in the integration of theta and HD processing. This convergence of signal processing early on, at the thalamic level, might be important to synchronize subcortical and cortical spatial activities (Korotkova et al., 2018), in turn facilitating (Aggleton and O’Mara, 2022) or maintaining downstream spatial computations in RSC via cortico-thalamo-cortical loops (Perry et al., 2021). For instance, according to the model by (Yan et al., 2021), landmark-responsive “multidirectional” HD cells in dRSC (Jacob et al., 2017; Zhang et al., 2022) receive thalamocortical inputs from Theta-by-HD cells. By firing on different theta phases, these cells time the feedforward and feedback transfer, between visual and entorhinal-hippocampal cortices via the dRSC. This coordinates visual landmark integration and dissociation, respectively, into the HD system.

This proposal is consistent with recent models of medial temporal lobe functions (Aggleton and O’Mara, 2022) suggesting that episodic memory relies on the oscillatory synchronization between two memory systems: hippocampal (place cells) and diencephalic (HD cells). This is thought to occur in shared parahippocampal areas, like the RSC, that receive inputs and whose activity is controlled by both streams (Aggleton and O’Mara, 2022; Chambers et al., 2021; Opalka et al., 2020; Yamawaki et al., 2019). In this case, AV is proposed to provide the coordination between cortical and subcortical spatial systems, via AV-RSC pathway(s), with theta providing the spatio-temporal organization for this transfer.

### Conclusion

Based on our observations, we have suggested a new role of AV, beyond its role as a theta relay to hippocampus: that is, the transfer (mostly via AVDM) and integration (mostly via AVVL) of theta and spatial signals (head orientation and spatial location). Overall, our results help to understand how the brain uses HD information to construct an increasingly complex cognitive map and shine a new light on AV as a building block of this map. We argued that the HD signal becomes integrated functionally into the hippocampal spatial code via AV-RSC pathway(s).

## Limitations of the study

Our study used tetrodes, which provide stable recordings but a low yield of cells, meaning that results needed to be compiled over a great number of animals and recording sessions. Additionally, because of the need to advance the electrodes between sessions, the precise location of each neuron could not be assessed directly but only reconstructed post hoc using microscopy together with documentation of the amount of electrode advancement.

However, the consistency of findings between animals compensates for this to some extent. Future work making use of new silicon probe high density recording technology will be able to replicate these findings with a larger data set.

We reported cells with locational selectivity that displayed place fields that were highly spatially localized, round and, in cases of multiple place fields, regularly arranged. Given the small size of our testing box (90 × 90cm), it may have been possible to miss-classify grid cells as place cells, as we would only be able to detect small grid scales (Savelli et al., 2008). While we have ruled out electrode-dragging issues and axonal recordings to explain our place cell results, evidence of synchronous firing in the spike cross-correlation between co-recorded place and Theta-by-HD cells would provide final indication that cells are monosynaptically connected. Although we did not find any evidence of such correlated activity, it is noteworthy that, even if place cells were part of the AV neuronal circuit rather than outsiders, chances of finding a place cell-non place cell pair that were co-recorded and also shared one synapse would be considerably low.

## Author contributions

E.L., A.S.M and K.J. designed the study. E.L. conducted the experiments and analyzed the data. A.S.M. assisted with supervision. E.L. and K. J. produced the visualizations. All authors wrote the paper and contributed to revision.

## Acknowledgments

The authors would like to thank Roddy M. Grieves for help with analysis and Brook Perry for assistance with experiments.

This work and A.S.M. were supported by grants from Wellcome Trust (WT 110157/Z/15/Z). K.J. was supported by grants from Wellcome Trust (103896AIA) and the Royal Society (IEC\R2\181140). E.L. was supported by a Medical Sciences Graduate School Studentship, University of Oxford: Clarendon Fund – Department of Experimental Psychology –Somerville College Mary Somerville Graduate Studentship (SFF1718_CB2_ MSD_1074636). For the purpose of open access, the authors have applied a CC BY public copyright to any Author Accepted Manuscript version arising from this submission.

## Tables with titles and legends

**Table S1:** Summary of recording locations and cell distribution in AV for all animals (n=733 cells in six animals). Related to Results: Histology, Fig. 12.

| Recorded cells |  |  |  |
| --- | --- | --- | --- |
| Rat | Hemisphere | Region | N |
| R222 | Left | AVDM | 0 |
|  |  | AVVL | 24 |
| R448 | Left | AVDM | 106 |
|  |  | AVVL | 76 |
| R449 | Left | AVDM | 0 |
|  |  | AVVL | 211 |
| R651 | Right | AVDM | 31 |
|  |  | AVVL | 66 |
| R652 | Left | AVDM | 116 |
|  |  | AVVL | 32 |
| R762 | Left | AVDM | 71 |
|  |  | AVVL | 0 |
| **All (n = 6)** |  | **AVDM** | **324** |
|  |  | **AVVL** | **409** |

**Table S2:** Numbers and percentages of AV cell types recorded in each animal. Related to Results: Multiple functional cell types in AV.

| Rat | N | Non-rhythmic HD | Theta-by-HD | Place | Theta | Unclassified |
| --- | --- | --- | --- | --- | --- | --- |
| R222 | 24 | 0 | 3 (13%) | 13 (54%) | 8 (33%) | 0 |
| R448 | 182 | 30 (17%) | 31 (17%) | 10 (6%) | 88 (48%) | 23 (13%) |
| R449 | 211 | 2 (1%) | 53 (25%) | 30 (14%) | 81 (38%) | 45 (21%) |
| R651 | 97 | 3 (3%) | 37 (38%) | 2 (2%) | 34 (35%) | 21 (22%) |
| R652 | 148 | 80 (54%) | 19 (13%) | 0 | 46 (31%) | 3 (2%) |
| R762 | 71 | 7 (10%) | 15 (24%) | 6 (9%) | 28 (39%) | 15 (21%) |
| **All** | **733** | **122 (17%)** | **158 (22%)** | **61 (8%)** | **285 (39%)** | **107 (15%)** |

**Table S3:** Electrophysiological properties of functional cell types in AV by subregion. Related to Results.

|  | Non-rhythmic HD | Theta-by-HD | Theta non-dir. | Place |
| --- | --- | --- | --- | --- |
| N cells |  |  |  |  |
| Total (/733) | 122 (17%) | 158 (22%) | 285 (39%) | 61 (8%) |
| AVDM (/324) | 69 (21%) | 53 (17%) | 146 (45%) | 17 (5%) |
| AVVL (/409) | 53 (13%) | 105 (26%) | 139 (34%) | 44 (11%) |
| Spike width (μs) | | | | |
| Total | 258.36 ± 13.73 | 267.98 ± 10.39 | 344.77 ± 9.70 | 284.26 ± 20.35 |
| AVDM | 258.26 ± 14.84 | 320.38 ± 19.1 | 366.03 ± 14.91 | 190.59 ± 16.81 |
| AVVL | 358.49 ± 25.21 | 241.52 ± 11.54 | 322.45 ± 12.03 | 320.45 ± 25.51 |
| Mean rate (Hz) |  |  |  |  |
| Total | 4.64 ± 0.42 | 2.55 ± 0.18 | 4.27 ± 0.32 | 1.1 ± 0.1 |
| AVDM | 3.45 ± 0.24 | 2.42 ± 0.29 | 4.70 ± 0.52 | 0.83 ± 0.17 |
| AVVL | 6.19 ± 0.74 | 2.62 ± 0.23 | 3.85 ± 0.38 | 1.20 ± 0.12 |
| Peak rate (Hz) |  |  |  |  |
| Total | 27.06 ± 2.92 | 5.20 ± 0.31 | 5.57 ± 0.38 | 11.29 ± 0.96 |
| AVDM | 19.57 ± 3.16 | 5.23 ± 0.36 | 6.07 ± 0.61 | 9.42 ± 1.86 |
| AVVL | 36.80 ± 5.04 | 5.16 ± 0.60 | 5.06 ± 0.44 | 12.01 ± 1.11 |
| Index of coupling (IC) |  |  |  |  |
| Total | 0.05 ± 0.00 | 0.17 ± 0.01 | 0.23 ± 0.01 | 0.15 ± 0.01 |
| AVDM | 0.05 ± 0.00 | 0.19 ± 0.02 | 0.25 ± 0.01 | 0.17 ± 0.02 |
| AVVL | 0.05 ± 0.01 | 0.16 ± 0.01 | 0.20 ± 0.01 | 0.14 ± 0.01 |
| Index of rhythmicity (IR) |  |  |  |  |
| Total | -0.004 ± 0.01 | 0.26 ± 0.01 | 0.24 ± 0.01 | 0.24 ± 0.02 |
| AVDM | 0.007 ± 0.02 | 0.26 ± 0.03 | 0.26 ± 0.02 | 0.24 ± 0.04 |
| AVVL | -0.02 ± 0.02 | 0.26 ± 0.02 | 0.20 ± 0.01 | 0.24 ± 0.03 |
| Bursting cell proportion |  |  |  |  |
| Total | 9/122 (7%) | 74/158 (47%) | 95/285 (33%) | 42/61 (69%) |
| AVDM | 5/69 (7%) | 24/53 (45%) | 46/146 (34%) | 13/17 (76%) |
| AVVL | 4/53 (8%) | 50/105 (48%) | 49/139 (35%) | 29/44 (66%) |
| Peak ISI (ms) |  |  |  |  |
| Total | 23.89 ± 3.16 | 10.24 ± 1.19 | 18.29 ± 1.89 | 5.75 ± 0.22 |
| AVDM | 31.72 ± 5.34 | 12.73 ± 2.88 | 18.67 ± 2.71 | 5.47 ± 0.32 |
| AVVL | 13.68 ± 1.11 | 8.98 ± 1.05 | 17.89 ± 2.64 | 5.86 ± 0.27 |
| Directional information (bits/spike) |  |  |  |  |
| Total | 1.10 ± 0.06 | 0.32 ± 0.02 | 0.05 ± 0.00 | 0.40 ± 0.04 |
| AVDM | 0.96 ± 0.08 | 0.34 ± 0.05 | 0.05 ± 0.00 | 0.58 ± 0.1 |
| AVVL | 1.28 ± 0.10 | 0.31 ± 0.02 | 0.05 ± 0.00 | 0.33 ± 0.04 |
| Spatial information. (bits/spike) |  |  |  |  |
| Total | 0.08 ± 0.01 | 0.31 ± 0.02 | 0.12 ± 0.01 | 1.37 ± 0.06 |
| AVDM | 0.09 ± 0.02 | 0.31 ± 0.04 | 0.10 ± 0.01 | 1.36 ± 0.11 |
| AVVL | 0.07 ± 0.01 | 0.31 ± 0.02 | 0.13 ± 0.02 | 1.38 ± 0.07 |
| Sparsity score |  |  |  |  |
| Total | 0.56 ± 0.01 | 0.49 ± 0.01 | 0.65 ± 0.01 | 0.15 ± 0.01 |
| AVDM | 0.55 ± 0.01 | 0.48 ± 0.02 | 0.65 ± 0.01 | 0.13 ± 0.01 |
| AVVL | 0.56 ± 0.01 | 0.49 ± 0.02 | 0.66 ± 0.01 | 0.16 ± 0.01 |
| Speed cell proportion |  |  |  |  |
| Total | 98/122 (80%) | 121/158 (77%) | 200/285 (70%) | 50/61 (82%) |
| AVDM | 55/69 (80%) | 38/53 (72%) | 99/146 (68%) | 12/17 (71%) |
| AVVL | 43/53 (81%) | 83/105 (79%) | 101/139 (73%) | 38/44 (87%) |
| Angular head velocity (AHV) cell proportion |  |  |  |  |
| Total | 11/122 (9%) | 24/158 (15%) | 0/285 | 9/61 (15%) |
| AVDM | 7/69 (10%) | 12/53 (23%) | 0/146 | 1/17(6%) |
| AVVL | 4/53 (8%) | 12/105 (11%) | 0/139 | 8/44 (18%) |

**Table S4:**
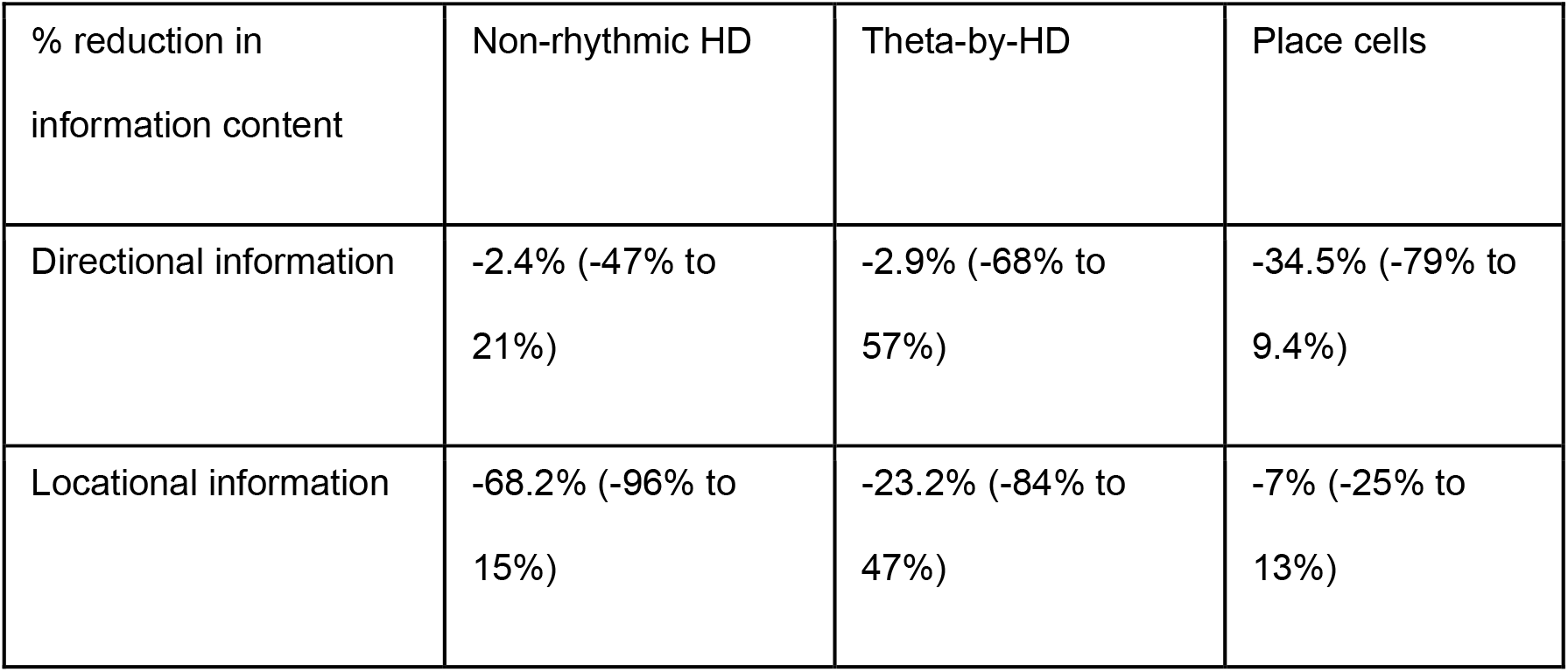
Percentage of reduction in locational and directional information in bits/spike after applying the pxd algorithm. Related to Supplementary information: Pxd-correction analysis for spatial information content.

| % reduction in information content | Non-rhythmic HD | Theta-by-HD | Place cells |
| --- | --- | --- | --- |
| Directional information | -2.4% (-47% to 21%) | -2.9% (-68% to 57%) | -34.5% (-79% to 9.4%) |
| Locational information | -68.2% (-96% to 15%) | -23.2% (-84% to 47%) | -7% (-25% to 13%) |

**Table S5:** Analysis of tuning curve shape for right (CW) and left (CCW) turns, and standard. Related to Supplementary Information: AHV had an effect on direction, but not tuning curve shape.

|  | CCW | CW | Standard |
| --- | --- | --- | --- |
| Tuning width (degrees) |  |  |  |
| Non-rhythmic HD | 93.29 ± 3.03 | 95.1 ± 2.99 | 92.08 ± 3.080 |
| Theta-by-HD | 131.53 ± 1.32 | 131.5 ± 1.23 | 130.81 ± 1.29 |
| Peak rate (Hz) |  |  |  |
| Non-rhythmic HD | 28.22 ± 3.05 | 28.57 ± 3.15 | 26.49 ± 2.91 |
| Theta-by-HD | 5.81 ± 0.38 | 5.96 ± 0.40 | 5.34 ± 0.35 |

## STAR Methods

### Animals

Six male Lister Hooded rats (Charles River, UK) weighing between 420 and 480g at time of surgery were used for electrophysiology recordings. Prior to the implant surgery, rats were handled for up to 10 days and familiarized with transportation to the recording lab and open field square arena. Animals were housed singly post-implantation in transparent Plexiglass cages in a temperature and humidity-controlled environment (21°C). Ten days after recovery from surgery, food regulation started (20g/day/rat) to maintain 85-90% of free-feeding body weight and testing began. All animals had free access to water throughout all experiments. Testing was conducted in the light phase of a 12h light/dark cycle (lights on at 7am). All experiments were carried out in accordance with UK Animals (Scientific Procedures) Act, 1986 and EU directive (2010-63-EU), complying with ARRIVE guidelines for the care and use of laboratory animals.

### Surgery

Standard stereotaxic surgery techniques for implantation of chronic recording electrodes were followed (Szymusiak and Nitz, 2003). Five animals were implanted in the left hemisphere and one in the right. Of these six animals, two were implanted in AVVL (R222 and R449) and four in AVDM (R762, R448, R651 and R652). In three of the AVDM animals (R448, R651 and R652), electrodes were advanced throughout the entire AV extent, sampling first AVDM and then moving into AVVL (see Table S1 for details on the cells recorded for all animals and AV subfields). AVDM implant coordinates were: AP, -1.7; ML, ± 1.4 − 1.5; DV, 3.6 (in mm from Bregma). AVVL implant coordinates were: AP, -1.7; ML, ± 1.7 − 1.8; DV, 3.4.

### Behavioral protocol and testing apparatus

Animals were screened daily for single unit activity. Screening and testing sessions took place inside a 90×90cm square arena with 60cm high walls, within a cue-rich room. The walls and floor of the box were covered with black vinyl sheeting. Animals were transferred inside a black box to the experimental room from their homecage, and connected to the multichannel recording apparatus (DacqUSB, Axona). Each trial was initiated via remote control after placing the rat inside the arena at a pseudo-random location and orientation.

Animals were screened daily for single unit activity. Screenings were made in an open field box within a cue-rich room. Spikes were monitored and when a single unit was isolated using TINT software (Axona, UK), the rat was removed from the arena and transferred back to homecage. The arena’s floor and walls were cleaned, and the rat was placed back in the arena for the experimental session.

Each basic experimental session was composed of two trials (“Light 1”, “Light 2” trials), lasting 8 or 16 minutes depending on the animal, during which rats foraged for rice scattered inside the arena. Animals were never disoriented prior to being placed in the arena. In this way, we did not disrupt internal sources of spatial information (Dudchenko et al., 1997). During each inter-trial interval, animals were removed from the arena and transferred back to the homecage, and the arena was cleaned to remove uncontrolled intra-maze non-visual cues. In some sessions, cells from four of the six implanted rats were additionally recorded in the same square box after switching off the room lights and switching on an infrared light (“Dark” trial), to assess the contribution of visual inputs to spatial firing patterns.

### Video tracking and signal collection/processing

During recording, the position and head direction of the animal were obtained from video tracking of two light-emitting diodes (LEDs) on the headstage, one large and one small and separated by 7cm. The LEDs were imaged with an overhead camera at a 50Hz sampling rate. Simultaneously, single unit and local field potential (LFP) signals were sampled at 48kHz using a headstage amplifier plugged into the microdrive connector and routed to the multichannel recording system (DacqUSB, Axona, UK) via a flexible lightweight cable. For single units, the signal from each electrode was differentially recorded against the signal from an electrode on a different tetrode with low spiking activity. It was amplified 15-50k times and band-pass filtered from 300Hz to 7kHz. For LFP, the signal was collected single-ended (250Hz), amplified 5K times, filtered with a 500Hz low-pass filter and a 50Hz notch filter.

Spike times, LFP signal, position in x- and y-coordinates and directional heading in degrees were saved for offline analysis.

### Data analysis

Data were analyzed offline using TINT (Axona Ltd) and MATLAB R2018a (MathWorks, USA) with custom-written software. The CircStat toolbox was used for circular statistics (Berens, 2009).

#### Spike sorting

Spike-sorting was performed using KlustaKwik (Kadir et al., 2014) followed by manual refinement using TINT. Single-unit inclusion criteria were (1) an average waveform displaying the classical action potential shape, and (2) fewer than 10% of spikes occurring in the first 2ms of the autocorrelogram (action potential refractory period). Only cells meeting these criteria were accepted into further analysis.

#### Waveform analysis

The width of the average spike waveform was used to separate narrow from broad spiking cell types (Lewicki, 1998) using a clustering protocol proposed by (Ardid et al., 2015)(open-source code from the public Git repository: https://bitbucket.org/sardid/waveformAnalysis). We fit the distribution of waveform widths with two Gaussian probability distributions. Two cutoffs were defined on these models as points at which the cumulative density function (CDF) for one Gaussian distribution was 10 times larger than the CDF for the other Gaussian i.e., the probability to belong to a group was 10 times larger than the probability to belong to the other one. Width values smaller than the first cutoff were classified as narrow, whereas values higher than the second cutoffss were classified as broad. Intermediate waveforms were those falling into the dip of the bimodal distribution. The calibrated version of the Hartigan Dip Test (<u>Hartigan and Hartigan 1985</u>) discarded unimodality for the distribution (*p* < 0.01).

#### Theta analysis

The Fast-Fourier transform (FFT) was used to find the LFP power spectrum. To compare across trials and animals, power measures within each trial were normalized to z-scores and plotted and smoothed with a Gaussian kernel (bandwidth = 2Hz; standard deviation = 0.5Hz). Theta frequency power is the maximum z-score within the 6-12 Hz theta frequency range. Average theta frequency is the frequency at peak theta power.

For phases analyses, the LFP signal was bandpass filtered at theta frequency range (6-12Hz) with a 4th order Butterworth filter. The Hilbert transform was applied to the bandpass-filtered LFP to derive instantaneous theta phases. Each theta cycle was defined such that peaks occurred at 0 and 360°, and troughs at 180°. To determine at which point to apply the Hilbert Transform, the LFP signal (Hz) was linearly interpolated with spike time (s), with each spike being assigned to the phase of the theta cycle at which it occurred. To visualize the spike-LFP relationship of a cell, firing phases were double-plotted in a frequency histogram. The probability distribution of spikes relative to theta phases was derived by smoothing the spike-phase histogram using a circular kernel density estimation (KDE) method, with an automatically selected bandwidth parameter using the plug-in rule of Taylor, 2008).

To obtain the relationship to running speed, the Hilbert transform was applied to the Butterworth filtered LFP signal to derive instantaneous theta frequency and amplitude that matched with running speed information recorded by the camera (50Hz). Instantaneous running speed was calculated as the distance between two consecutive position points, divided by the time between them. To analyze the relationship between instantaneous theta power vs running speed, we binned speed data into 10cm/s bin, within a range of 0-50cm/s. Average theta power was computed for each bin, and a linear regression model was fitted to the data using the least-squares approach. The slope of the regression line and Pearson’s linear correlation coefficient (r) between speed bins and the corresponding power were derived. The same method was used to analyze the relationship between instantaneous theta frequency and speed.

Spike-LFP coherence and rhythmicity were assessed by computing an index of rhythmicity (IR) and an index of theta phase-coupling (IC). Rhythmicity refers to the frequency of the temporal modulation of the spike-time autocorrelogram at theta frequency range (6-12Hz), while coupling refers to the phase of theta at which spikes occurred. We used both measures because spikes may be emitted with a timing phase-locked to theta even if there is no overt autocorrelogram rhythmicity (Alexander et al., 2020; Eliav et al., 2018).

Autocorrelograms of the spike trains were plotted between ± 500ms using 10ms bins, normalized to the maximum value and smoothed (20 bins boxcar). The IR was calculated as the difference between the expected theta modulation-trough (autocorrelogram value between 60-70ms) and the theta-modulation peak (autocorrelogram value between 120-130ms), divided by their sum. It takes values between -1 and 1 (Lozano et al., 2017). Cells were considered as theta modulated if they passed the 99^th^ percentile shuffle cutoff for IC and had an IR>=0.001 (Tsanov et al., 2010, 2011). For phase-locked cells, the preferred theta phase was found as the circular mean of all the spike phases. For the IC, phases were binned between 0 and 360° with 6° bins and the IC was found as the mean vector length (Rayleigh vector, CircStat toolbox) of these angles (Climer et al., 2015; Frank et al., 2001).

Units that fire locked to theta oscillations can fire on every theta cycle, or on alternate ones (theta skipping; Brandon et al., 2013). To derive the theta skip index (TS), a curve was fitted to the autocorrelogram (10ms bins between ± 400ms) using the equation in Brandon et al., 2013:

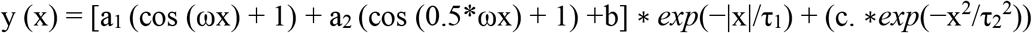

with x=autocorrelogram bins, m=maximum autocorrelation value. Fit parameters were restricted to the following values: a_1_ = [0, m], a_2_ = [0, m], b = [0, m], c = [−m, m], ω = [10π, 18π], τ_1_ = [0, 5] and τ_2_ = [0, 0.05]. TS index, bound between -1 and 1, represents the difference between the height of the first and second peaks of the fitted curve, divided by the largest of the two. For this analysis, we selected only cells that passed our theta phase-locking criterion (IC >= 99^th^ percentile shuffling). Among these cells, a theta-skipping cell had to meet following criteria: (a) a good (R2 > 0.7) fit of the model parameters to the autocorrelogram; (b) a baseline theta power component in the autocorrelogram (IR >= 0.001), and (c) the first side peak of the autocorrelogram smaller than the second side peak (TS > 0.1; Brandon et al., 2013). All cells were confirmed by visual inspection of their autocorrelogram.

The approach used to investigate phase precession consisted of comparing the intrinsic oscillation of cells (theta frequency in the temporal autocorrelogram) to the global theta oscillation frequency (theta frequency in the LFP). When cells phase precess, they fire progressively earlier with each successive theta cycle, so on average the theta frequency in their autocorrelogram is slightly faster than the global LFP frequency (Meer and Redish, 2011). To derive intrinsic theta frequency, a decomposing sine wave of frequency ω was then fitted to the autocorrelogram (10ms bins between ± 500ms) using the equation in Grieves et al., 2020:

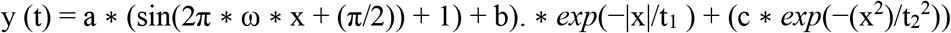

where t is the autocorrelogram time lag, and a-c, ω, and t_1-2_ were fit to the data using a non-linear least squares method (Matlab function *fit*). Parameters were restricted to the following values: ω = [6, 12], a-b = [0 Inf], c = [0, 0.8], t_1_-_2_ = [0, 0.05]. Intrinsic theta frequency corresponds to the fit parameter, ω (Royer et al., 2010).

#### Directional tuning curves

Directionality in the firing of a cell was quantified by the mean vector length (R-vector) of the directional tuning curve. Spike and HD data were sorted into 60 bins of 6°. The mean firing rates per angular bin (Hz) were smoothed with a 5-bin (30°) smoothing kernel and polar-plotted. The parameters that were used to quantify tuning curve characteristics were the mean vector length, or Rayleigh vector (R-vector, CircStat toolbox), directional peak firing rate (Hz), PFD (directional bin associated with the peak firing rate), tuning width (two standard deviations from the circular mean direction (φ), and concentration parameter, *k* (Zar, 2010). Cells were considered as directionally modulated if they passed the 99^th^ percentile shuffle cutoff for R-vector. Directional information content in bits/spike was also computed following the method of Skaggs et al., 1992, which computes the amount of spatial information carried by each spike.

To analyze drift, we quantified the amount of variability in the firing direction of a cell when it fired >50% of its peak firing rate for the trial. We took all spikes emitted at these times (from the spike-time histogram), plus spikes emitted before and after it up to an angular distance of +/-50°. This defines a full sweep of the head through the cell’s PFD. A selection of spikes based on each individual sweep rather than on a reference frame from the whole dataset meant that we did not need the PFD prior, which would bias our sampling of spikes in case of drift. For each full-sweep event, we computed the circular KDE of the HD, using an automatically selected bandwidth parameter k (Taylor, 2008). The KDE method was used to avoid arbitrary decisions on binning and because distributional properties are more easily assessed compared to a histogram (assessed next). Since the reliability of the density estimate depends on the number of data points (Fisher, 1993), only sweeps containing at least 5 spikes were considered.

For each trial, we found the average width of all the KDE peaks (width at half-height), and how much variability there was in the absolute direction of these peaks (SEM of peaks location) and compared between cell groups using two-sample T-tests. Doing so allowed us to test two different, but non-mutually exclusive, hypotheses to explain broader tuning curves of Theta-by-HD cells: (1) tuning curves are inherently broader due to the directional firing range actually being larger, or (2) they reflect a less stable directional signal, due to PFD shifting to a larger extent during the trial. If the former is correct, there should be little jitter in the KDE peaks (small SEM of peaks location) but average width of the KDEs should be large. If the latter is correct, there should be a larger variability in KDE peaks (larger SEM of KDEs location) but small mean peak width.

#### Self-motion correlates

We quantified the relationship between instantaneous firing rates and linear/angular speed of the head using a speed score (s-score) and an AHV score, respectively. The instantaneous firing rate of a cell was derived from the smoothed spike-time histogram, obtained using 20ms bins (coinciding to 50Hz sampling rate), smoothed with a 250ms-wide Gaussian kernel. Dividing the spike count per bin by 0.02 converts counts to firing rate for each position data point. Instantaneous running speed was measured as the distance between two consecutive position data points, divided by the time between them. Instantaneous firing rate was binned by running speed (2cm/s bins, between 2 and 50cm/s) and fitted by a linear regression model using the least-squares approach. The s-score is the Pearson’s linear correlation coefficient between instantaneous firing rate and speed bins (Kropff et al., 2015). Speed-tuned cells were considered to be those with an absolute s-score above 0.3.

Instantaneous AHV was measured as the angular difference between two successive HD data points, divided by the time between them (Blair and Sharp, 1995). HD measurements were interpolated between consecutive samples to provide an estimate of HD at each AHV point. A 2°/s bin width was used to bin instantaneous firing rate by AHV, between -100 and 100°/s was used. A scattergram of firing rate vs AHV was plotted, and linear regression lines were fitted to AHV values between -2 and -50°/s, and between 2 and 50°/s. Values between -2°/s and 2°/s were discarded, as periods in which the head was not turning. CW and CCW AHV scores are the Pearson’s correlation between instantaneous firing rate and AHV bins. To quantify AHV modulation irrespective of the turning direction, we first considered absolute AHV values. AHV-responsive cells have an absolute AHV score above 0.3. Taking the absolute values was necessary because HD cells can both increase and decrease firing rates with AHV in different turning directions (Bassett and Taube, 2001; Lozano et al., 2017), having slopes that are positive in one direction and negative in the other, which cancel if raw values are taken. We then considered CW and CCW AHV scores separately to classify cells as symmetric or asymmetric.

#### Directional tuning curve cross-correlation

To compare cross-trial stability between non-rhythmic HD and Theta-by-HD cells, we used a cross-correlation approach. Smoothed tuning curves for each pair of trial (Light-Light; Light-Dark) were circularly cross-correlated in 6° steps and Pearson’s correlation (r) was computed at each step. Correlation at 0° was used as a measure of cross-trial stability for each cell. The distribution of all correlations was compared to the distribution of correlations derived from cross correlating tuning curves in the first trial with shuffled tuning curves from the second trial (spikes time shifted 10,000 times relative to HD). The 95^th^ percentile value from the distribution of shuffled correlations was taken as the chance-level criteria against which to test the similarity of directional fields for each cell.

#### Spatial analysis

Locational ratemaps were calculated by binning position and spikes data in 2×2cm bins, smoothed separately using a weighted Gaussian kernel (smoothing factor = 5cm) such that data points closer to a bin’s center had more influence on that bin’s firing rate. Bins that were >10cm away from the current bin had no influence (weight=0) and were not included in the Gaussian smoothing process. The smoothed ratemap was generated by dividing the smoothed spikemap by the smoothed dwelltime map (Grieves et al., 2020). The overall spike count divided by session duration gave the mean firing rate (Hz) of the cell. The locational peak firing rate (Hz) is the firing rate of the cell in the bin with the maximum average rate To correct for the sampling bias of the animal when it explores the box (which could lead to artefactual spatial selectivity), we used a maximum likelihood approach, known as the position-by-direction (pxd) correction (Burgess et al., 2005; Cacucci et al., 2004). Similar numbers of bins were applied to locations and directions. There were 60 6° directional bins and 64 locational bins. No smoothing was applied. This provides better convergence of the pxd algorithm and ensures that direction and location were given equal weighting in finding the solution. This method was applied to all cells (n=1 non-rhythmic HD cell for which the algorithm did not converge upon a solution was discarded).

Corrected measures of locational and directional information content and sparsity were calculated according to the formulas in (Skaggs et al., 1992). Thalamic units were classified as place cells based on high locational information and a low sparsity, computed as the percentage of the arena’s surface over which a cell fired. A place field was defined as a group of at least 9 adjoining pixels in which firing rate was at least 20% of the peak firing rate, and peak firing rate was at least 1Hz.

#### Cell classification

To generate control data, a shuffling procedure was used in which each cell’s spike train was circularly time-shifted 10,000 times by a random amount between 20s and the duration of the recording session minus 20s, relative to the behavioral or theta phase data. The relevant scores were recomputed each time and values were pooled together to derive a control distribution for theta and HD modulation. A 99^th^ percentile cutoff was used to determine significance levels for individual cells to be theta- or HD-modulated. Given that the R-vector measure is very sensitive to small amounts of unimodality, setting the threshold high ensured that only cells with clear theta and HD selectivity made it into the sample and that the classified cells would certainly be considered as theta or HD had they been recorded from a different brain region.

The shuffle procedure was then used for cell classification, in addition to a number of other criteria as follows. A cell was considered theta-modulated if it fired rhythmic spike trains (IR >= 0.001) locked to theta (IC >= 99^th^ shuffle percentile). A non-rhythmic HD cell was defined as a cell with significant directionality (directional peak firing rate > 1Hz; R-vector >= 99^th^ shuffle percentile) that was not also theta-modulated. A Theta-by-HD cell showed both theta and HD modulation. A place cell was a cell with significant locational modulation (mean firing rate > 0.1Hz but < 10Hz; locational peak firing rate > 1Hz; locational information content > 0.8 bits/spike; sparsity score < 0.2). All the place cells reported were also theta-modulated. Conservative thresholds were chosen intentionally to set a lower bound for the proportions of place cells in the AV and ensure that only units with high locational selectivity were included in the sample.

#### Cell-specific temporal firing characteristics

To investigate temporal firing properties, inter-spike interval (ISI) histograms were created by binning all ISIs in the range 0-250ms with 2ms bins. The peak ISI of a cell was taken as the centre of the histogram bin with the highest count. A cell was classified as bursting if peak ISI < 6ms. The burst index was defined as the ratio of spikes sharing ISI < 6ms to all spikes emitted by the cell in a trial.

#### Anticipatory time interval (ATI) analysis

The anticipatory time interval (ATI) was quantified using two methods. In the first method, the ATI of a cell was estimated based on cross-correlating clockwise (CW) and counter-clockwise (CCW) tuning functions for that cell (Blair and Sharp, 1995). For each cell, CW and CCW spikes were identified according to the angular head velocity (AHV) at which the head was turning when a spike occurred. CW spikes occurred at AHV > 60°/s, CCW spikes occurred at AHV < -60°/s. Spikes emitted at times in which the head was not turning were removed. CW and CCW spikes were time-shifted in steps of 20ms, from 20 to 160ms (coinciding with 50Hz sampling frequency). For each step, CW and CCW tuning curves were plotted, normalized to 1Hz and circularly cross-correlated by rotating one relative to the other in steps of 6° to find the rotation angle that produced the maximal cross-correlation (Pearson’s r) between the two curves. Angles were plotted as a function of the corresponding time shift, and a linear regression model was fitted using the least squares method. The ATI is the time shift required to align CW and CCW functions. The difference angle is the rotation angle that yields the maximal cross-correlation between CCW and CW tuning functions for a time shift of 0ms. Cells with cross-correlation below 0.7 were removed (non-rhythmic HD, n=10; Theta-by-HD, n=30). In the second method, the cell’s ATI was derived based on optimal time-shift calculations for the standard tuning function. A cell’s optimal shift corresponds to the amount of time shifts required to optimize its tuning curve parameters. Optimal shifts were computed by shifting the entire spikes-time series (CW and CCW spikes combined) forward and backward in time in steps of 20ms, between 20 and 160ms, and between -20 and -160ms, and plotting the tuning curve at each step to derive peak rate and tuning width. The second method was used because cross-correlations of tuning functions might not provide a robust ATI estimate for cells with tuning curves that are not perfectly symmetric, such as those that have a higher dispersion and multiple peaks, as those seen here for Theta-by-HD cells.

### Histology

After completion of all recordings, rats were anesthetized with isoflurane and received an overdose of sodium pentobarbital (Euthatal, Merial, Harlow, UK). Rats were perfused transcardially using saline (0.9% sodium chloride solution) followed by 10% formalin perfusion (10% formalin in 0.9% sodium chloride solution). The brains were removed and post fixed overnight in 10% formalin, transferred to a 25-30% sucrose solution (25-30% in 0.1M PBS) for a minimum of 48h for cryoprotection. Brains were sliced coronally into sections of 40μm thickness on a freezing microtome. Sections were mounted on glass slides.

Tetrode locations were histologically verified from cresyl violet stained brain sections. Photomicrographs were captured using a camera mounted on a Leica DMR microscope (Leica Microsystems, UK) with a x1.6 objective.

### Statistics

If assumption of normality was met, comparisons between two groups were conducted using independent-samples or paired-samples t-tests. Otherwise, we used non-parametric statistical tests and comparisons between groups were conducted using the two-sample Wilcoxon Rank-sum (WRS) tests and paired-sample Wilcoxon Signed rank (WSR) tests. For more than two groups, a Kruskal-Wallis (KW) test was conducted, followed by multiple pairwise comparisons of group medians (SPSS, Bonferroni correction). A Chi-square test tested differences in the observed number of cells between two groups against an expected equal proportion. Two-sample Kolmogorov-Smirnov test (KS test) tested whether two cumulative density functions (cdf) differed significantly.

For circular data, a Watson Williams (WW) multi-sample F-test compared circular means between two or more groups. Equality of the two circular distributions was tested using a Kuiper test (Fisher, 1993), and MATLAB functions *circ_ktest* for differences in concentration parameters. All statistical tests are two-tailed. In all figures and tables, *p*-value significance level: *=0.05 significance level, **=0.01, ***=0.001.

## References

Aggleton, J.P., Brown, M.W., 1999. Episodic memory, amnesia, and the hippocampal-anterior thalamic axis. Behav. Brain Sci. 22, 425–489.

Aggleton, J.P., Hunt, P.R., Nagle, S., Neave, N., 1996. The effects of selective lesions within the anterior thalamic nuclei on spatial memory in the rat. Behav. Brain Res. 81, 189– 198. https://doi.org/10.1016/s0166-4328(96)89080-2

Aggleton, J.P., O’Mara, S.M., 2022. The anterior thalamic nuclei: core components of a tripartite episodic memory system. Nat. Rev. Neurosci. 23, 505–516. https://doi.org/10.1038/s41583-022-00591-8

Aggleton, J.P., O’Mara, S.M., Vann, S.D., Wright, N.F., Tsanov, M., Erichsen, J.T., 2010. Hippocampal–anterior thalamic pathways for memory: uncovering a network of direct and indirect actions. Eur. J. Neurosci. 31, 2292–2307. https://doi.org/10.1111/j.1460-9568.2010.07251.x

Albo, Z., Viana Di Prisco, G., Vertes, R.P., 2003. Anterior thalamic unit discharge profiles and coherence with hippocampal theta rhythm. Thalamus Relat. Syst. 2, 133–144. https://doi.org/10.1016/S1472-9288(03)00006-2

Alexander, A.S., Carstensen, L.C., Hinman, J.R., Raudies, F., Chapman, G.W., Hasselmo, M.E., 2020. Egocentric boundary vector tuning of the retrosplenial cortex. Sci. Adv. 6, eaaz2322. https://doi.org/10.1126/sciadv.aaz2322

Alexander, A.S., Nitz, D.A., 2015. Retrosplenial cortex maps the conjunction of internal and external spaces. Nat. Neurosci. 18, 1143–1151. https://doi.org/10.1038/nn.4058

Ardid, S., Vinck, M., Kaping, D., Marquez, S., Everling, S., Womelsdorf, T., 2015. Mapping of Functionally Characterized Cell Classes onto Canonical Circuit Operations in Primate Prefrontal Cortex. J. Neurosci. 35, 2975–2991. https://doi.org/10.1523/JNEUROSCI.2700-14.2015

Bassett, J.P., Taube, J.S., 2001. Neural Correlates for Angular Head Velocity in the Rat Dorsal Tegmental Nucleus. J. Neurosci. 21, 5740–5751. https://doi.org/10.1523/JNEUROSCI.21-15-05740.2001

Berens, P., 2009. CircStat: A MATLAB Toolbox for Circular Statistics. J. Stat. Softw. 31, 1– 21. https://doi.org/10.18637/jss.v031.i10

Blair, H.T., Sharp, P.E., 1995. Anticipatory head direction signals in anterior thalamus: evidence for a thalamocortical circuit that integrates angular head motion to compute head direction. J. Neurosci. 15, 6260–6270.

Brandon, M.P., Bogaard, A.R., Schultheiss, N.W., Hasselmo, M.E., 2013. Segregation of cortical head direction cell assemblies on alternating theta cycles. Nat. Neurosci. 16, 739–748. https://doi.org/10.1038/nn.3383

Burgess, N., Cacucci, F., Lever, C., O’keefe, J., 2005. Characterizing multiple independent behavioral correlates of cell firing in freely moving animals. Hippocampus 15, 149– 153. https://doi.org/10.1002/hipo.20058

Cacucci, F., Lever, C., Wills, T.J., Burgess, N., O’Keefe, J., 2004. Theta-modulated place-by-direction cells in the hippocampal formation in the rat. J. Neurosci. 24, 8265–8277. https://doi.org/10.1523/JNEUROSCI.2635-04.2004

Calton, J.L., Stackman, R.W., Goodridge, J.P., Archey, W.B., Dudchenko, P.A., Taube, J.S., 2003. Hippocampal Place Cell Instability after Lesions of the Head Direction Cell Network. J. Neurosci. 23, 9719–9731. https://doi.org/10.1523/JNEUROSCI.23-30-09719.2003

Casali, G., Bush, D., Jeffery, K., 2019. Altered neural odometry in the vertical dimension. Proc. Natl. Acad. Sci. 116, 4631–4636. https://doi.org/10.1073/pnas.1811867116

Chambers, A.R., Berge, C.N., Vervaeke, K., 2021. Cell-type-specific silence in thalamocortical circuits precedes hippocampal sharp-wave ripples. bioRxiv 2021.05.05.442741. https://doi.org/10.1101/2021.05.05.442741

Christiansen, K., Dillingham, C.M., Wright, N.F., Saunders, R.C., Vann, S.D., Aggleton, J.P., 2016. Complementary subicular pathways to the anterior thalamic nuclei and mammillary bodies in the rat and macaque monkey brain. Eur. J. Neurosci. 43, 1044–1061. https://doi.org/10.1111/ejn.13208

Climer, J.R., DiTullio, R., Newman, E.L., Hasselmo, M.E., Eden, U.T., 2015. Examination of rhythmicity of extracellularly recorded neurons in the entorhinal cortex. Hippocampus 25, 460–473. https://doi.org/10.1002/hipo.22383

Csicsvari, J., Hirase, H., Czurkó, A., Mamiya, A., Buzsáki, G., 1999. Oscillatory Coupling of Hippocampal Pyramidal Cells and Interneurons in the Behaving Rat. J. Neurosci. 19, 274–287. https://doi.org/10.1523/JNEUROSCI.19-01-00274.1999

Dudchenko, P.A., Goodridge, J.P., Taube, J.S., 1997. The effects of disorientation on visual landmark control of head direction cell orientation. Exp. Brain Res. 115, 375–380. https://doi.org/10.1007/pl00005707

Eliav, T., Geva-Sagiv, M., Yartsev, M.M., Finkelstein, A., Rubin, A., Las, L., Ulanovsky, N., 2018. Nonoscillatory Phase Coding and Synchronization in the Bat Hippocampal Formation. Cell 175, 1119-1130.e15. https://doi.org/10.1016/j.cell.2018.09.017

Fenton, A.A., Kao, H.-Y., Neymotin, S.A., Olypher, A., Vayntrub, Y., Lytton, W.W., Ludvig, N., 2008. Unmasking the CA1 Ensemble Place Code by Exposures to Small and Large Environments: More Place Cells and Multiple, Irregularly Arranged, and Expanded Place Fields in the Larger Space. J. Neurosci. 28, 11250–11262. https://doi.org/10.1523/JNEUROSCI.2862-08.2008

Fisher, N.I., 1993. Statistical Analysis of Circular Data. Cambridge University Press, Cambridge. https://doi.org/10.1017/CBO9780511564345

Frank, L.M., Brown, E.N., Wilson, M.A., 2001. A Comparison of the Firing Properties of Putative Excitatory and Inhibitory Neurons From CA1 and the Entorhinal Cortex. J. Neurophysiol. 86, 2029–2040. https://doi.org/10.1152/jn.2001.86.4.2029

Frost, B.E., Martin, S.K., Cafalchio, M., Islam, M.N., Aggleton, J.P., O’Mara, S.M., 2021. Anterior Thalamic Inputs Are Required for Subiculum Spatial Coding, with Associated Consequences for Hippocampal Spatial Memory. J. Neurosci. 41, 6511– 6525. https://doi.org/10.1523/JNEUROSCI.2868-20.2021

Gabriel, M., 1993. Discriminative Avoidance Learning: A Model System, in: Vogt, B.A., Gabriel, M. (Eds.), Neurobiology of Cingulate Cortex and Limbic Thalamus: A Comprehensive Handbook. Birkhäuser, Boston, MA, pp. 478–523. https://doi.org/10.1007/978-1-4899-6704-6_18

Goodridge, J.P., Taube, J.S., 1997. Interaction between the postsubiculum and anterior thalamus in the generation of head direction cell activity. J. Neurosci. 17, 9315–9330.

Grieves, R.M., Jedidi-Ayoub, S., Mishchanchuk, K., Liu, A., Renaudineau, S., Jeffery, K.J., 2020. The place-cell representation of volumetric space in rats. Nat. Commun. 11, 789. https://doi.org/10.1038/s41467-020-14611-7

Harding, A., Halliday, G., Caine, D., Kril, J., 2000. Degeneration of anterior thalamic nuclei differentiates alcoholics with amnesia. Brain J. Neurol. 123 (Pt 1), 141–154. https://doi.org/10.1093/brain/123.1.141

Harland, B., Contreras, M., Souder, M., Fellous, J.-M., 2021. Dorsal CA1 hippocampal place cells form a multi-scale representation of megaspace. Curr. Biol. CB 31, 2178-2190.e6. https://doi.org/10.1016/j.cub.2021.03.003

Jacob, P.-Y., Casali, G., Spieser, L., Page, H., Overington, D., Jeffery, K., 2017. An independent, landmark-dominated head direction signal in dysgranular retrosplenial cortex. Nat. Neurosci. 20, 173–175. https://doi.org/10.1038/nn.4465

Jankowski, M.M., Islam, M.N., Wright, N.F., Vann, S.D., Erichsen, J.T., Aggleton, J.P., O’Mara, S.M., 2014. Nucleus reuniens of the thalamus contains head direction cells. eLife 3, e03075. https://doi.org/10.7554/eLife.03075

Jankowski, M.M., O’Mara, S.M., 2015. Dynamics of place, boundary and object encoding in rat anterior claustrum. Front. Behav. Neurosci. 9. https://doi.org/10.3389/fnbeh.2015.00250

Jankowski, M.M., Passecker, J., Islam, M.N., Vann, S., Erichsen, J.T., Aggleton, J.P., O’Mara, S.M., 2015. Evidence for spatially-responsive neurons in the rostral thalamus. Front. Behav. Neurosci. 9. https://doi.org/10.3389/fnbeh.2015.00256

Jankowski, M.M., Ronnqvist, K.C., Tsanov, M., Vann, S.D., Wright, N.F., Erichsen, J.T., Aggleton, J.P., O’Mara, S.M., 2013. The anterior thalamus provides a subcortical circuit supporting memory and spatial navigation. Front. Syst. Neurosci. 7, 45. https://doi.org/10.3389/fnsys.2013.00045

Kadir, S.N., Goodman, D.F.M., Harris, K.D., 2014. High-dimensional cluster analysis with the Masked EM Algorithm. Neural Comput. 26, 2379–2394. https://doi.org/10.1162/NECO_a_00661

King, C., Recce, M., O’Keefe, J., 1998. The rhythmicity of cells of the medial septum/diagonal band of Broca in the awake freely moving rat: relationships with behaviour and hippocampal theta. Eur. J. Neurosci. 10, 464–477. https://doi.org/10.1046/j.1460-9568.1998.00026.x

Kocsis, B., Di Prisco, G.V., Vertes, R.P., 2001. Theta synchronization in the limbic system: the role of Gudden’s tegmental nuclei. Eur. J. Neurosci. 13, 381–388.

Korotkova, T., Ponomarenko, A., Monaghan, C.K., Poulter, S.L., Cacucci, F., Wills, T., Hasselmo, M.E., Lever, C., 2018. Reconciling the different faces of hippocampal theta: The role of theta oscillations in cognitive, emotional and innate behaviors. Neurosci. Biobehav. Rev. 85, 65–80. https://doi.org/10.1016/j.neubiorev.2017.09.004

Kropff, E., Carmichael, J.E., Moser, M.-B., Moser, E.I., 2015. Speed cells in the medial entorhinal cortex. Nature 523, 419–424. https://doi.org/10.1038/nature14622

Leutgeb, S., Ragozzino, K.E., Mizumori, S.J.Y., 2000. Convergence of head direction and place information in the CA1 region of hippocampus. Neuroscience 100, 11–19. https://doi.org/10.1016/S0306-4522(00)00258-X

Lever, C., Burton, S., Jeewajee, A., O’Keefe, J., Burgess, N., 2009. Boundary Vector Cells in the Subiculum of the Hippocampal Formation. J. Neurosci. 29, 9771–9777. https://doi.org/10.1523/JNEUROSCI.1319-09.2009

Lewicki, M.S., 1998. A review of methods for spike sorting: the detection and classification of neural action potentials. Netw. Bristol Engl. 9, R53–78.

Lomi, E., Mathiasen, M.L., Cheng, H.Y., Zhang, N., Aggleton, J.P., Mitchell, A.S., Jeffery, K.J., 2021. Evidence for two distinct thalamocortical circuits in retrosplenial cortex. Neurobiol. Learn. Mem. 185, 107525. https://doi.org/10.1016/j.nlm.2021.107525

Lozano, Y.R., Page, H., Jacob, P.-Y., Lomi, E., Street, J., Jeffery, K., 2017. Retrosplenial and postsubicular head direction cells compared during visual landmark discrimination. Brain Neurosci. Adv. 1. https://doi.org/10.1177/2398212817721859

Meer, M.A.A. van der, Redish, A.D., 2011. Theta Phase Precession in Rat Ventral Striatum Links Place and Reward Information. J. Neurosci. 31, 2843–2854. https://doi.org/10.1523/JNEUROSCI.4869-10.2011

Mizuseki, K., Buzsáki, G., 2013. Preconfigured, skewed distribution of firing rates in the hippocampus and entorhinal cortex. Cell Rep. 4, 1010–1021. https://doi.org/10.1016/j.celrep.2013.07.039

Mizuseki, K., Royer, S., Diba, K., Buzsáki, G., 2012. Activity Dynamics and Behavioral Correlates of CA3 and CA1 Hippocampal Pyramidal Neurons. Hippocampus 22, 1659–1680. https://doi.org/10.1002/hipo.22002

O’Keefe, J., Recce, M.L., 1993. Phase relationship between hippocampal place units and the EEG theta rhythm. Hippocampus 3, 317–330. https://doi.org/10.1002/hipo.450030307

Opalka, A.N., Huang, W., Liu, J., Liang, H., Wang, D.V., 2020. Hippocampal Ripple Coordinates Retrosplenial Inhibitory Neurons during Slow-Wave Sleep. Cell Rep. 30, 432-441.e3. https://doi.org/10.1016/j.celrep.2019.12.038

Paxinos, G., Watson, C., 2007. The Rat Brain in Stereotaxic Coordinates: Hard Cover Edition, 6th edition. ed. Academic Press, Amsterdam ; Boston.

Perry, B.A.L., Lomi, E., Mitchell, A.S., 2021. Thalamocortical interactions in cognition and disease: The mediodorsal and anterior thalamic nuclei. Neurosci. Biobehav. Rev. 130, 162–177. https://doi.org/10.1016/j.neubiorev.2021.05.032

Perry, B.A.L., Mitchell, A.S., 2019. Considering the Evidence for Anterior and Laterodorsal Thalamic Nuclei as Higher Order Relays to Cortex. Front. Mol. Neurosci. 12, 167. https://doi.org/10.3389/fnmol.2019.00167

Poulter, S., Lee, S.A., Dachtler, J., Wills, T.J., Lever, C., 2021. Vector trace cells in the subiculum of the hippocampal formation. Nat. Neurosci. 24, 266–275. https://doi.org/10.1038/s41593-020-00761-w

Royer, S., Sirota, A., Patel, J., Buzsáki, G., 2010. Distinct Representations and Theta Dynamics in Dorsal and Ventral Hippocampus. J. Neurosci. 30, 1777–1787. https://doi.org/10.1523/JNEUROSCI.4681-09.2010

Savelli, F., Yoganarasimha, D., Knierim, J.J., 2008. Influence of boundary removal on the spatial representations of the medial entorhinal cortex. Hippocampus 18, 1270–1282. https://doi.org/10.1002/hipo.20511

Seki, M., Zyo, K., 1984. Anterior thalamic afferents from the mamillary body and the limbic cortex in the rat. J. Comp. Neurol. 229, 242–256. https://doi.org/10.1002/cne.902290209

Sharp, P.E., 1999. Complimentary roles for hippocampal versus subicular/entorhinal place cells in coding place, context, and events. Hippocampus 9, 432–443. https://doi.org/10.1002/(SICI)1098-1063(1999)9:4<432::AID-HIPO9>3.0.CO;2-P

Sharp, P.E., Green, C., 1994. Spatial correlates of firing patterns of single cells in the subiculum of the freely moving rat. J. Neurosci. 14, 2339–2356.

Shibata, H., 1994. Terminal distribution of projections from the retrosplenial area to the retrohippocampal region in the rat, as studied by anterograde transport of biotinylated dextran amine. Neurosci. Res. 20, 331–336. https://doi.org/10.1016/0168-0102(94)90055-8

Shibata, H., 1993a. Direct projections from the anterior thalamic nuclei to the retrohippocampal region in the rat. J. Comp. Neurol. 337, 431–445. https://doi.org/10.1002/cne.903370307

Shibata, H., 1993b. Efferent projections from the anterior thalamic nuclei to the cingulate cortex in the rat. J. Comp. Neurol. 330, 533–542. https://doi.org/10.1002/cne.903300409

Shibata, H., 1992. Topographic organization of subcortical projections to the anterior thalamic nuclei in the rat. J. Comp. Neurol. 323, 117–127. https://doi.org/10.1002/cne.903230110

Shibata, H., 1989. Descending projections to the mammillary nuclei in the rat, as studied by retrograde and anterograde transport of wheat germ agglutinin-horseradish peroxidase. J. Comp. Neurol. 285, 436–452. https://doi.org/10.1002/cne.902850403

Shibata, H., Yoshiko, H., 2015. Thalamocortical projections of the anteroventral thalamic nucleus in the rabbit. J. Comp. Neurol. 523, 726–741. https://doi.org/10.1002/cne.23700

Skaggs, W., McNaughton, B., Gothard, K., 1992. An Information-Theoretic Approach to Deciphering the Hippocampal Code, in: Advances in Neural Information Processing Systems. Morgan-Kaufmann.

Szymusiak, R., Nitz, D., 2003. Chronic recording of extracellular neuronal activity in behaving animals. Curr. Protoc. Neurosci. 6, 6.16. https://doi.org/10.1002/0471142301.ns0616s21

Taube, J.S., 1995. Head direction cells recorded in the anterior thalamic nuclei of freely moving rats. J. Neurosci. 15, 70–86.

Taube, J.S., Muller, R.U., Ranck, J.B., 1990. Head-direction cells recorded from the postsubiculum in freely moving rats. I. Description and quantitative analysis. J. Neurosci. 10, 420–435.

Taylor, C.C., 2008. Automatic bandwidth selection for circular density estimation. Comput. Stat. Data Anal. 52, 3493–3500. https://doi.org/10.1016/j.csda.2007.11.003

Tsanov, M., Chah, E., Vann, S.D., Reilly, R.B., Erichsen, J.T., Aggleton, J.P., O’Mara, S.M., 2011. Theta-Modulated Head Direction Cells in the Rat Anterior Thalamus. J. Neurosci. 31, 9489–9502. https://doi.org/10.1523/JNEUROSCI.0353-11.2011

Tsanov, M., Chah, E., Wright, N., Vann, S.D., Reilly, R., Erichsen, J.T., Aggleton, J.P., O’Mara, S.M., 2010. Oscillatory Entrainment of Thalamic Neurons by Theta Rhythm in Freely Moving Rats. J. Neurophysiol. 105, 4–17. https://doi.org/10.1152/jn.00771.2010

Tsivilis, D., Vann, S.D., Denby, C., Roberts, N., Mayes, A.R., Montaldi, D., Aggleton, J.P., 2008. A disproportionate role for the fornix and mammillary bodies in recall versus recognition memory. Nat. Neurosci. 11, 834–842. https://doi.org/10.1038/nn.2149

Van Groen, T., Wyss, J.M., 2003. Connections of the retrosplenial granular b cortex in the rat. J. Comp. Neurol. 463, 249–263. https://doi.org/10.1002/cne.10757

Van Groen, T., Wyss, J.M., 1995. Projections from the anterodorsal and anteroventral nucleus of the thalamus to the limbic cortex in the rat. J. Comp. Neurol. 358, 584– 604. https://doi.org/10.1002/cne.903580411

van Groen, T., Wyss, J.M., 1992. Connections of the retrosplenial dysgranular cortex in the rat. J. Comp. Neurol. 315, 200–216. https://doi.org/10.1002/cne.903150207

van Groen, T., Wyss, J.M., 1990a. Connections of the retrosplenial granular a cortex in the rat. J. Comp. Neurol. 300, 593–606. https://doi.org/10.1002/cne.903000412

van Groen, T., Wyss, J.M., 1990b. The postsubicular cortex in the rat: characterization of the fourth region of the subicular cortex and its connections. Brain Res. 529, 165–177. https://doi.org/10.1016/0006-8993(90)90824-u

van Groen, Thomas, Wyss, J.M., 1990. The connections of presubiculum and parasubiculum in the rat. Brain Res. 518, 227–243. https://doi.org/10.1016/0006-8993(90)90976-I

van Strien, N.M., Cappaert, N.L.M., Witter, M.P., 2009. The anatomy of memory: an interactive overview of the parahippocampal-hippocampal network. Nat. Rev. Neurosci. 10, 272–282. https://doi.org/10.1038/nrn2614

Vertes, R.P., Albo, Z., Viana Di Prisco, G., 2001. Theta-rhythmically firing neurons in the anterior thalamus: implications for mnemonic functions of Papez’s circuit. Neuroscience 104, 619–625. https://doi.org/10.1016/s0306-4522(01)00131-2

Vertes, R.P., Hoover, W.B., Viana Di Prisco, G., 2004. Theta rhythm of the hippocampus: subcortical control and functional significance. Behav. Cogn. Neurosci. Rev. 3, 173– 200. https://doi.org/10.1177/1534582304273594

Vertes, R.P., Kocsis, B., 1997. Brainstem-diencephalo-septohippocampal systems controlling the theta rhythm of the hippocampus. Neuroscience 81, 893–926. https://doi.org/10.1016/s0306-4522(97)00239-x

Warburton, E.C., Baird, A.L., Morgan, A., Muir, J.L., Aggleton, J.P., 2000. Disconnecting hippocampal projections to the anterior thalamus produces deficits on tests of spatial memory in rats. Eur. J. Neurosci. 12, 1714–1726. https://doi.org/10.1046/j.1460-9568.2000.00039.x

Welday, A.C., Shlifer, I.G., Bloom, M.L., Zhang, K., Blair, H.T., 2011. Cosine Directional Tuning of Theta Cell Burst Frequencies: Evidence for Spatial Coding by Oscillatory Interference. J. Neurosci. 31, 16157–16176. https://doi.org/10.1523/JNEUROSCI.0712-11.2011

Wilent, W.B., Nitz, D.A., 2007. Discrete Place Fields of Hippocampal Formation Interneurons. J. Neurophysiol. 97, 4152–4161. https://doi.org/10.1152/jn.01200.2006

Winter, S., Taube, J., 2014. Head Direction Cells: From Generation to Integration, in: Space, Time and Memory in the Hippocampal Formation. p. 24. https://doi.org/10.1007/978-3-7091-1292-2_4

Winter, S.S., Clark, B.J., Taube, J.S., 2015. Spatial navigation. Disruption of the head direction cell network impairs the parahippocampal grid cell signal. Science 347, 870–874. https://doi.org/10.1126/science.1259591

Witter, M.P., Ostendorf, R.H., Groenewegen, H.J., 1990. Heterogeneity in the Dorsal Subiculum of the Rat. Distinct Neuronal Zones Project to Different Cortical and Subcortical Targets. Eur. J. Neurosci. 2, 718–725. https://doi.org/10.1111/j.1460-9568.1990.tb00462.x

Wright, N.F., Erichsen, J.T., Vann, S.D., O’Mara, S., Aggleton, J.P., 2010. Parallel but separate inputs from limbic cortices to the mammillary bodies and anterior thalamic nuclei in the rat. J. Comp. Neurol. 518, 2334–2354. https://doi.org/10.1002/cne.22336

Yamawaki, N., Li, X., Lambot, L., Ren, L.Y., Radulovic, J., Shepherd, G.M.G., 2019. Long-range inhibitory intersection of a retrosplenial thalamocortical circuit by apical tuft-targeting CA1 neurons. Nat. Neurosci. 22, 618–626. https://doi.org/10.1038/s41593-019-0355-x

Yan, Y., Burgess, N., Bicanski, A., 2021. A model of head direction and landmark coding in complex environments. PLOS Comput. Biol. 17, e1009434. https://doi.org/10.1371/journal.pcbi.1009434

Yoder, R.M., Peck, J.R., Taube, J.S., 2015. Visual Landmark Information Gains Control of the Head Direction Signal at the Lateral Mammillary Nuclei. J. Neurosci. 35, 1354– 1367. https://doi.org/10.1523/JNEUROSCI.1418-14.2015

Zar, J.H., 2010. Biostatistical Analysis. Prentice Hall.

Zhang, N., Grieves, R.M., Jeffery, K.J., 2022. Environment Symmetry Drives a Multidirectional Code in Rat Retrosplenial Cortex. J. Neurosci. 42, 9227–9241. https://doi.org/10.1523/JNEUROSCI.0619-22.2022

Zhang, S., Schönfeld, F., Wiskott, L., Manahan-Vaughan, D., 2014. Spatial representations of place cells in darkness are supported by path integration and border information. Front. Behav. Neurosci. 8. https://doi.org/10.3389/fnbeh.2014.00222

