## Supplementary Information for "Convergence of direction, location and theta in the rat anteroventral thalamic nucleus"

### Related to Results: Multiple functional cell types in AV

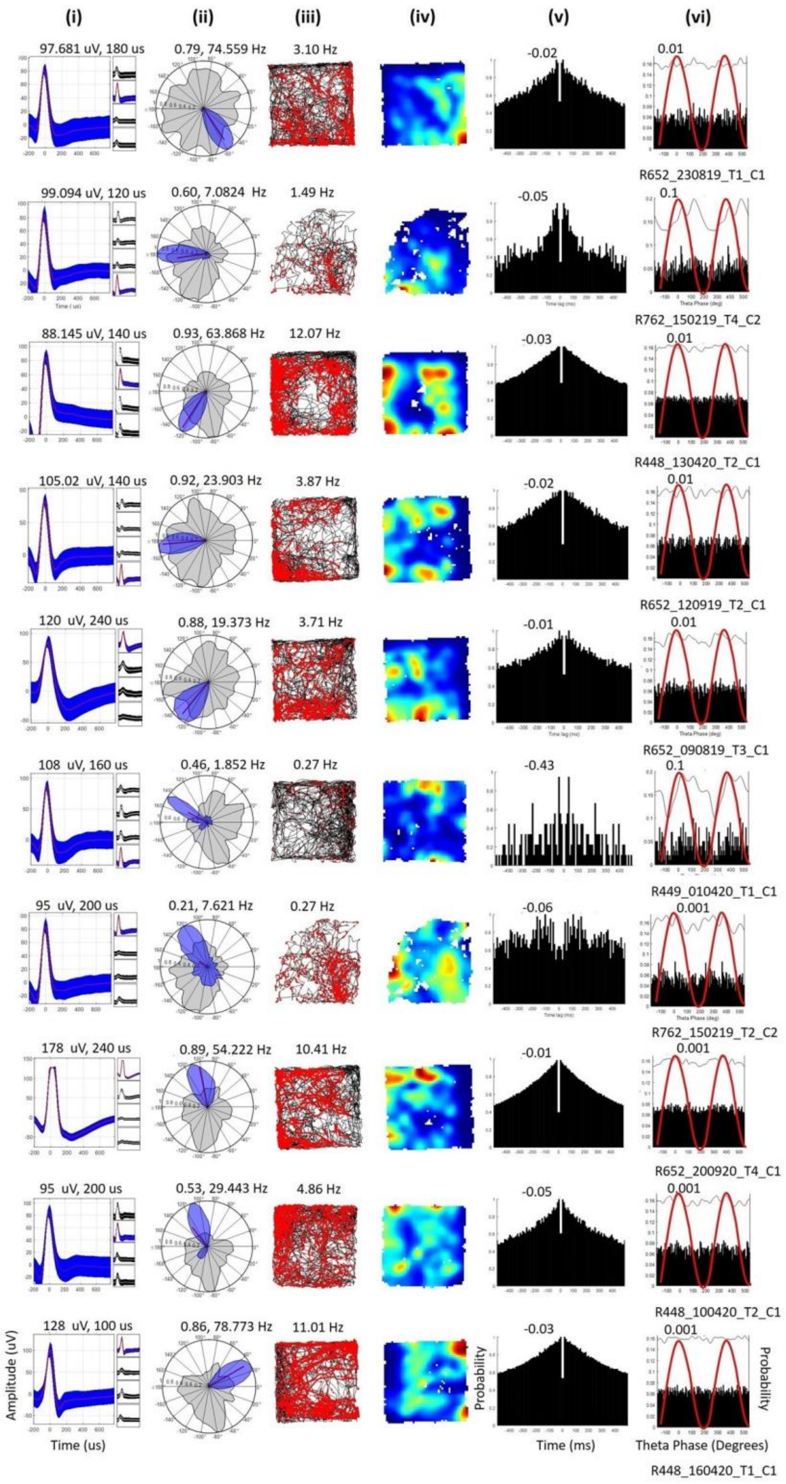


**Figure S1: Non-rhythmic HD cells. Related to Fig. 2A and displayed the same way.** Note the mix of broad and narrow waveforms despite similar directional tuning and theta modulation. Cells are tightly tuned to HD but lack locational selectivity and their firing is not theta-modulated.

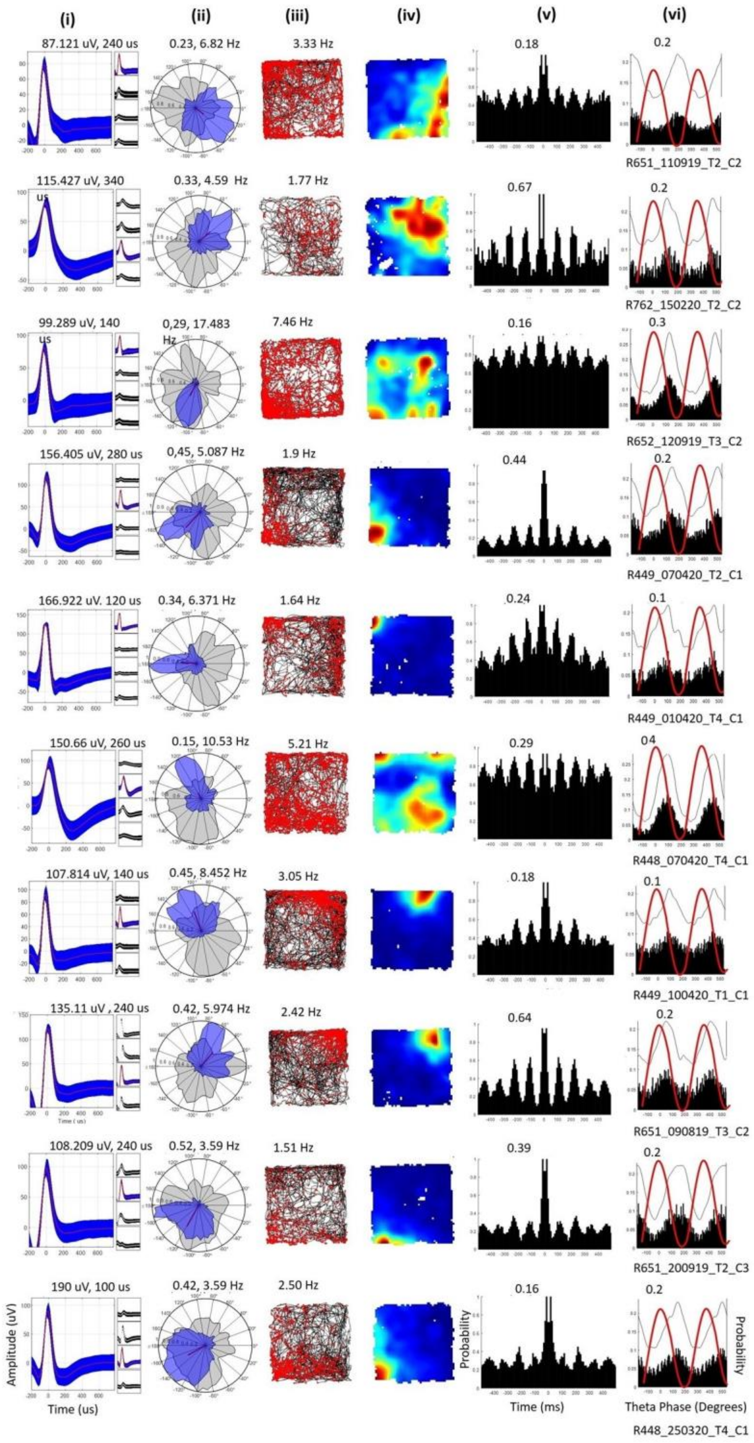


**Figure S2: Theta-by-HD cells, displayed as for Fig. 2A.** Note the mix of waveform types but broader directional tuning than for classic HD cells, and clear theta modulation. Some cells display broad patches of spatially inhomogeneous firing, in the form of broad firing fields in the ratemaps. The cells also show rhythmic, phase-locked spiking, aligned to descending theta phases, near theta troughs.

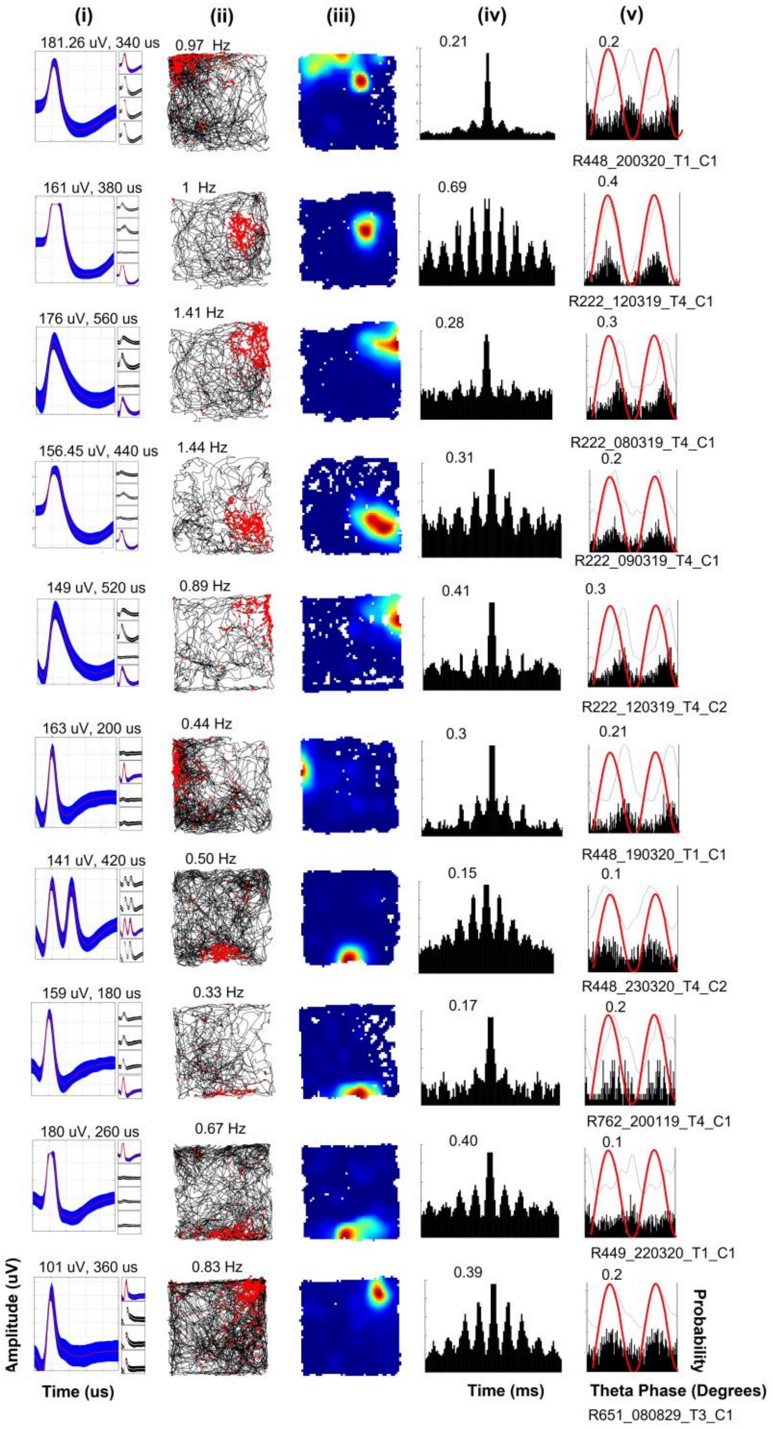


**Figure S3: Place cells, displayed as for Fig. 8A, with one single firing field.** Cells are rhythmic and phase-locked to descending theta phases, after theta peaks. Autocorrelograms show a high central peak (0 ms), suggestive of complex spike bursting.

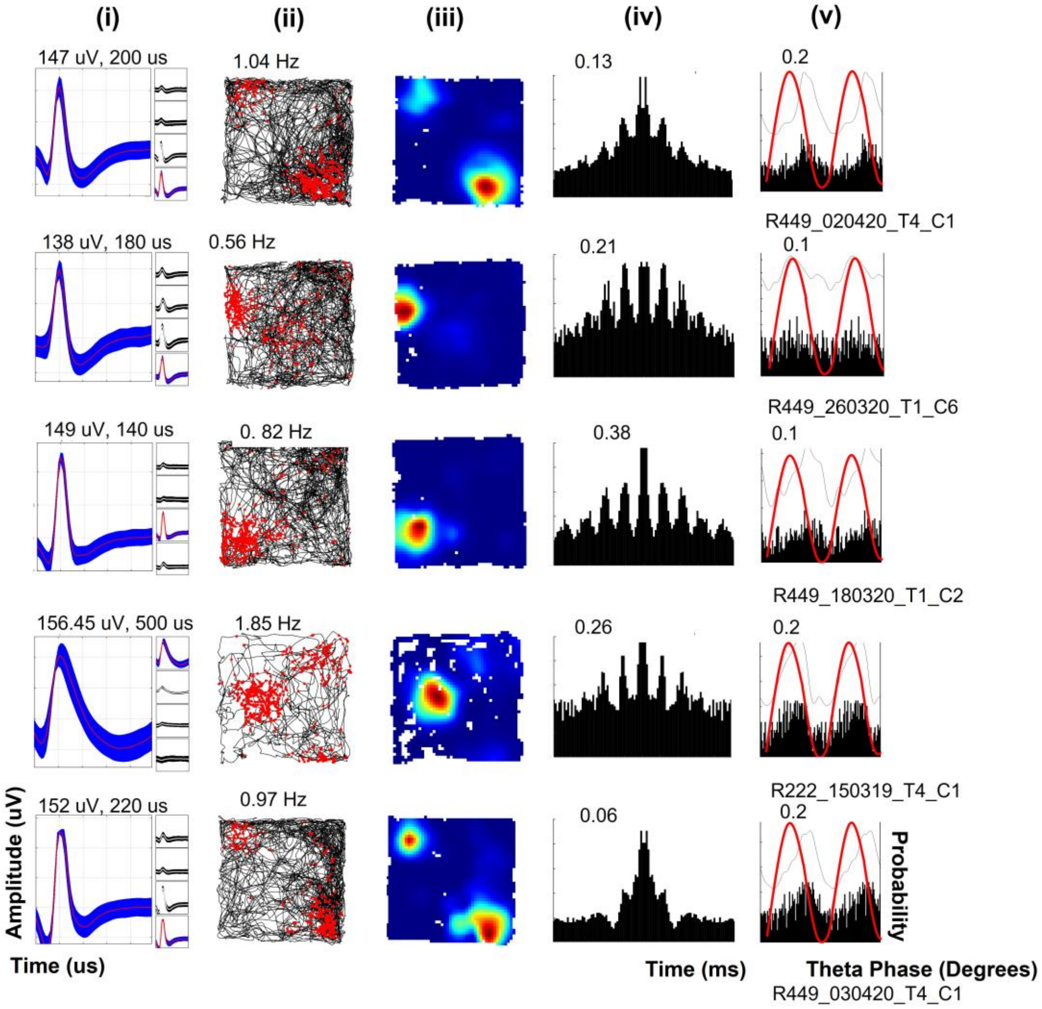


**Figure S4: Place cells, displayed as for Fig. 8A, with two firing fields (n = 5/61 cases).** Same caption as Fig. S3.

**Temporal firing properties. Related to Results: Waveform analysis reveals different neuronal subtypes.**

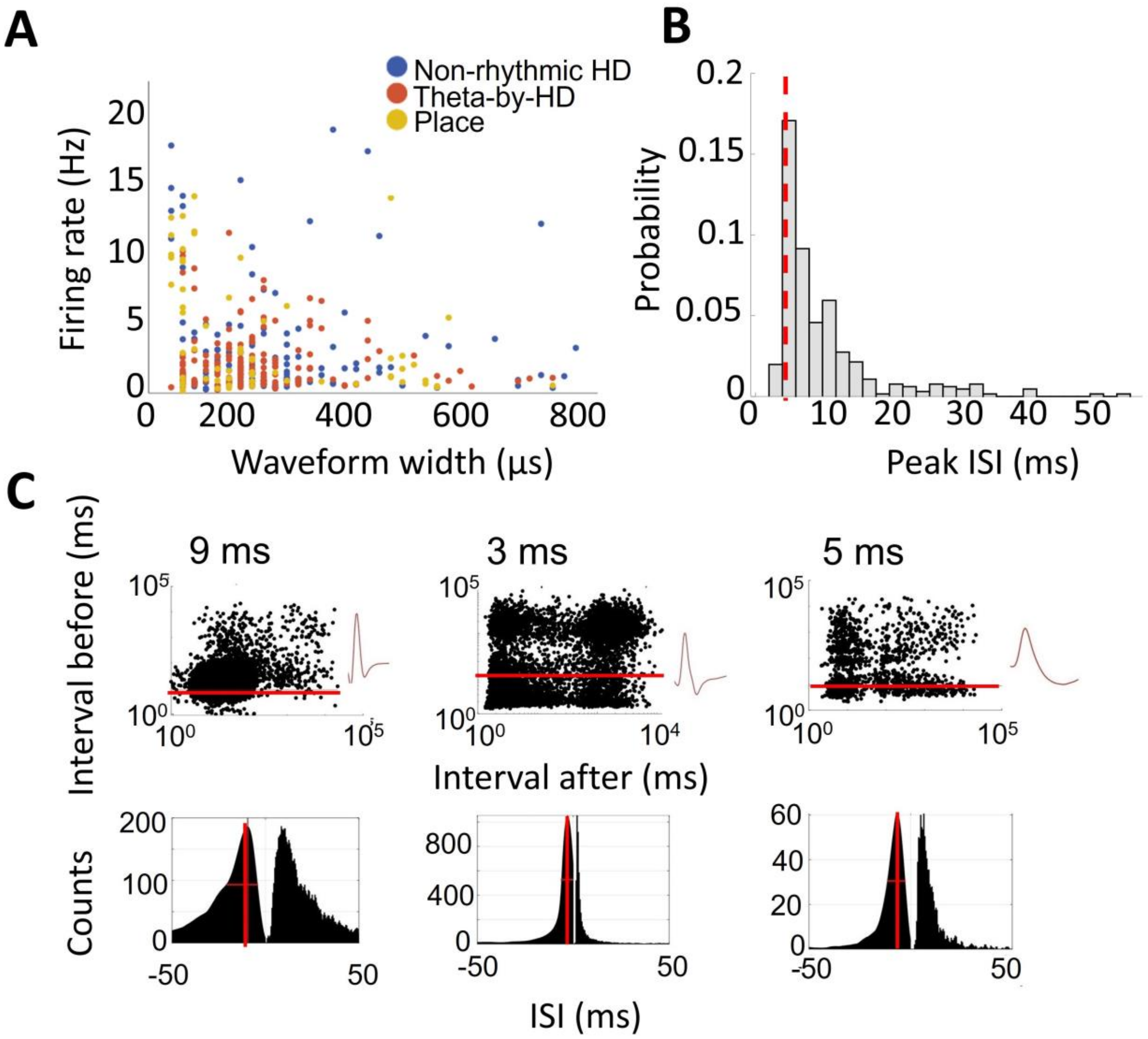


**Figure S5: Basic characterization of temporal firing properties. Related to Fig. 3C-D**. (A) Scatterplot of mean rate (Hz) vs peak-to-trough waveform width (μs) for the spatial cell types identified. Note variable waveform widths. (B) Probability distribution of peak inter-spike intervals (ISI) for the cells in A. Red line = 6ms. (C) ISI profiles of three example cells. Top, Log-scale ISI scatterplots. Each point is a spike, x-axis, interval before, y-axis, interval after. Red line marks the 6ms pre-ISI. Bottom, ISI histogram shown as raw and as smooth density estimate. Red line = 6ms pre-ISI. Theta-rhythmic and bursting neurons have a fast-decaying histogram peak and three ISI clusters, formed by spikes with an ISI of 120ms before or after them and inter-train spikes. In some cases, a fourth cluster is visible, formed by single spikes emitted at 6-12 frequencies (not part of a train). Average spike waveform is plotted next to each cell.

**Theta analysis. Related to Results: Oscillatory entrainment of unit activity by the local LFP theta**

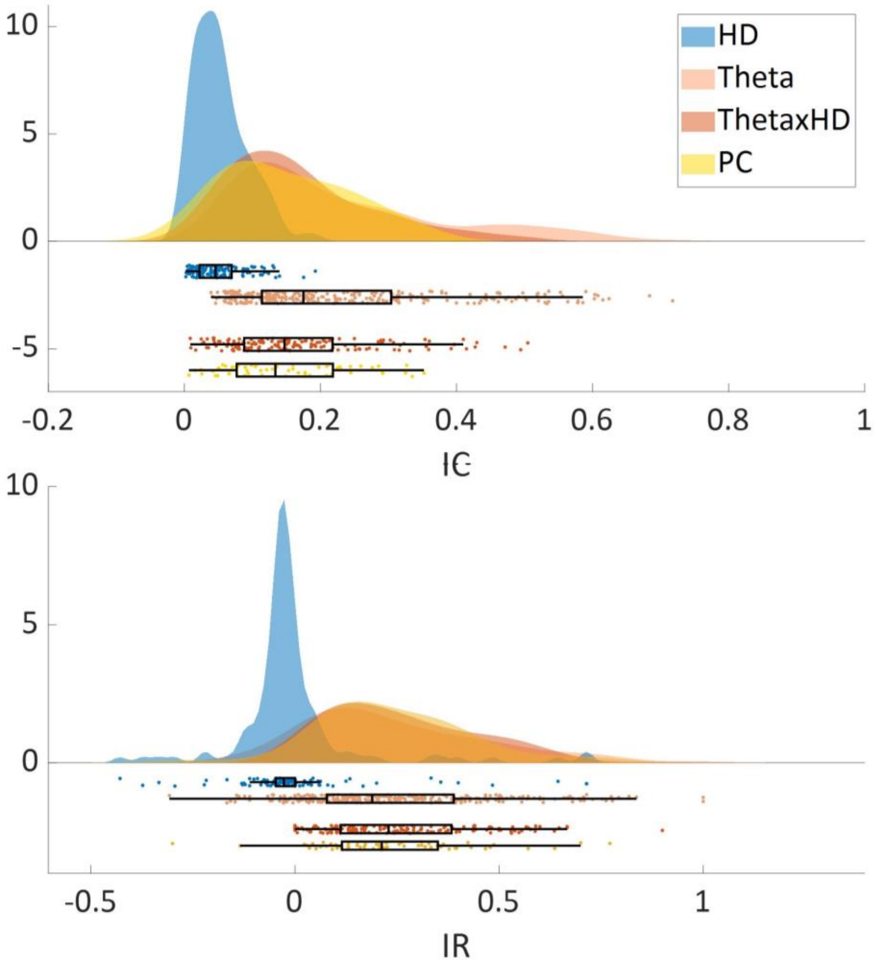


**Figure S6: Differences in theta-modulation, related to Fig. 5.** Distribution of index of phase coupling (IC) and index of rhythmicity (IR). Data is shown as a split violin plot + raw jittered data point + boxplot. Split violin plots show the probability density function (PDF) of the observed data points. Each marker is a cell. IC and IR did not differ between theta (non-HD), theta/rhythmic HD and place cells. All these cells fire locked to theta and show stronger theta-modulation depth compared to non-theta rhythmic HD cells (in blue).

A subset of Theta-by-HD cells (n = 46/158; 29.11%) showed evidence of theta cycle-skipping in their temporal autocorrelogram. These cells, named theta-skipping HD cells, mostly fired on alternating cycles, with only some spikes emitted at theta frequency (some cycles were not skipped), as indicated by the first autocorrelogram side-peak at 125ms being smaller than the second side-peak at 250ms (Fig. S7). Note that our cell-type inclusion criteria for Theta-by-HD cells based on IC and IR did not allow us to formally distinguish theta phase-locked units that were theta-rhythmic from phase-locked units that were rhythmic and skipping, given that they would display equal levels of locking and rhythmicity strength. To classify these cells, we derived a theta-skip index (TS) from the autocorrelogram that indicated how much the first side-peak was skipped by the firing activity. Among all 158 Theta-by-HD cells, 119 (75.3%) displayed a good autocorrelogram model fit (R2 > 0.7) and 47 of these cells (39.5%) exceeded the threshold for theta-skipping (TS > 0.1). One cell was removed by visual inspection, leaving a total of 46 theta-skipping HD cells. A total of 53/61 (86.9%) place cells displayed a good model fit and seven of these cells (13.7%) exceeded the threshold for theta-skipping. Six of these cells were later removed by visual inspection (variable autocorrelogram shape due to small spike number), leaving only one theta-skipping place cell that was not analysed further. None of the non-rhythmic HD cells met the requirement for a baseline theta power component in the autocorrelogram and thus were not considered for TS calculations. We found no difference in preferred theta phase, spatial nor temporal firing properties between Theta-by-HD cells that were rhythmic and Theta-by-HD cells that were rhythmic + skipping (not shown), except for differences in TS index magnitude (0.19 ± 0.01 vs -0.01 ± 0.01; two-sample WRS test, z = 9.28, $p < 0.0001$).

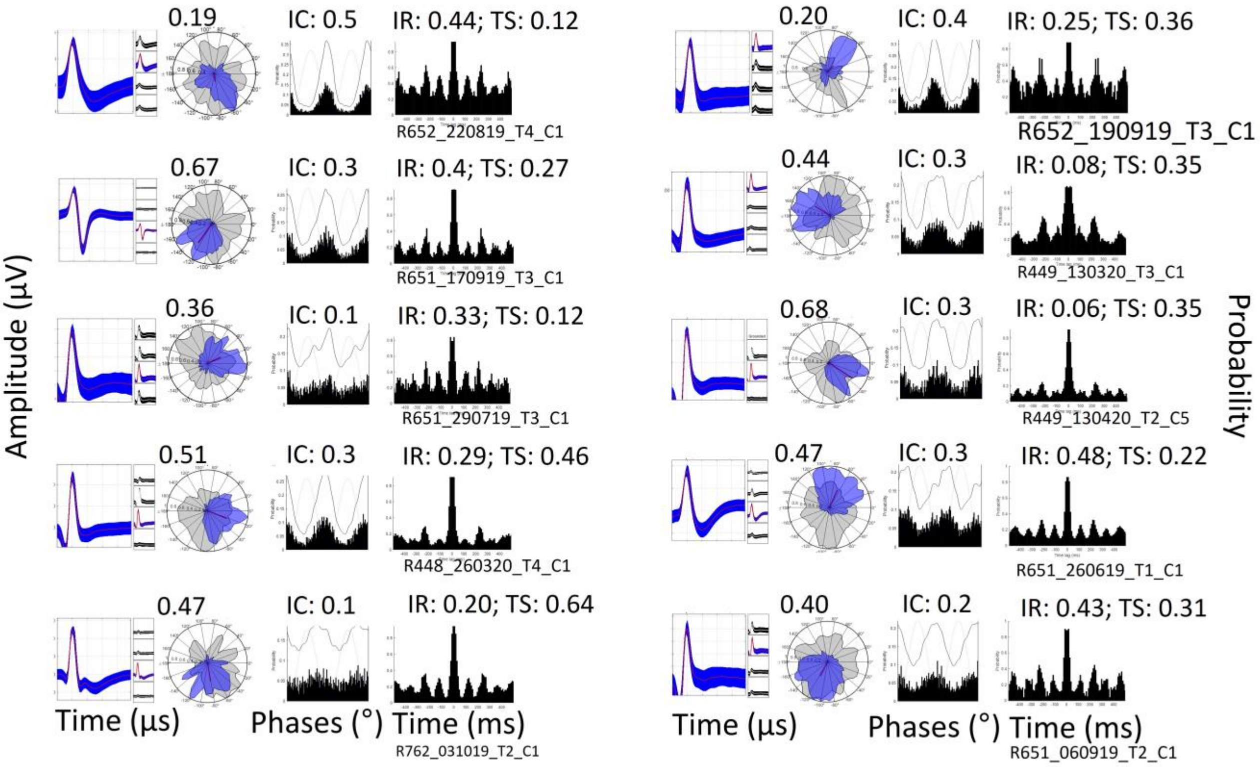


**Figure S7: Examples of theta-skipping HD cells.** For each cell, from left to right: average spike waveform; x-axis, time (µs); y-axis, amplitude (µV). Polar plot of directional tuning curve (blue) and dwelltime curve (gray), with R-vector. Double-plotted spikes time vs theta phases histogram, with Index of Coupling (IC). Peak-normalized temporal autocorrelogram implemented between ± 500ms, with Index of Rhythmicity (IR) and theta-skip index (TS). Cells fire locked to alternating descending theta phases (see Methods Theta analysis).

### Pxd-correction analysis for spatial information content. Related to Results: Spatial firing

To check that locational selectivity was not a sampling artefact or a by-product of directional firing, corrected measures of spatial information content (bits/spike) were computed (see STAR Methods: Spatial analysis). We used the code taken from Burgess et al., 2005, available online from these authors. This analysis confirmed that place cells were strongly locational modulated, non-rhythmic HD cells were strongly directional, and Theta-by-HD cells were intermediate for both measures. See main text results: 13% of Theta-by-HD cells displayed conjunctive tuning for place and orientation, in the form of broad directional and locational fields (example in Fig. S9).

Percentages of reduction of locational and directional information content after pxd correction are reported in Table S4 for each of group. Non-rhythmic HD cells had largely reduced locational but not directional information, while the opposite was true for place cells. Compared to non-rhythmic HD cells, Theta-by-HD cells showed a lower average reduction in both directional and locational information content, suggesting that coding of location was not due to their directional selectivity. Note the large range values, meaning that the specific effect of pxd correction on a given Theta-by-HD cell was highly variable. Fig. S8-S9 illustrate the effects of applying the pxd correction algorithm on two pairs of simultaneously recorded Theta-by-HD and place cells.

At the population level, average Pearson's correlations for direction were as follows: place cell, 0.77; non-rhythmic HD, 0.99; Theta-by-HD, 0.93. Average correlations for location were as follows: place cell, 0.997; non-rhythmic HD, 0.88; Theta-by-HD, 0.99. Correlation coefficients compared using Fishers r-to-z transformation were all statistically different from each other (all $p < 0.0001$). Specifically, Theta-by-HD cells were intermediate non-rhythmic HD and place cells for both location and direction correlations. Note that place cell correlations matched values reported for CA1 place cells (direction, 0.74; location, 0.96), while non-rhythmic HD cell correlation matched values for canonical HD cells in cortical regions (direction, 0.99; location, 0.48). Values were obtained using the same pxd approach as in the present work (Cacucci et al., 2004).

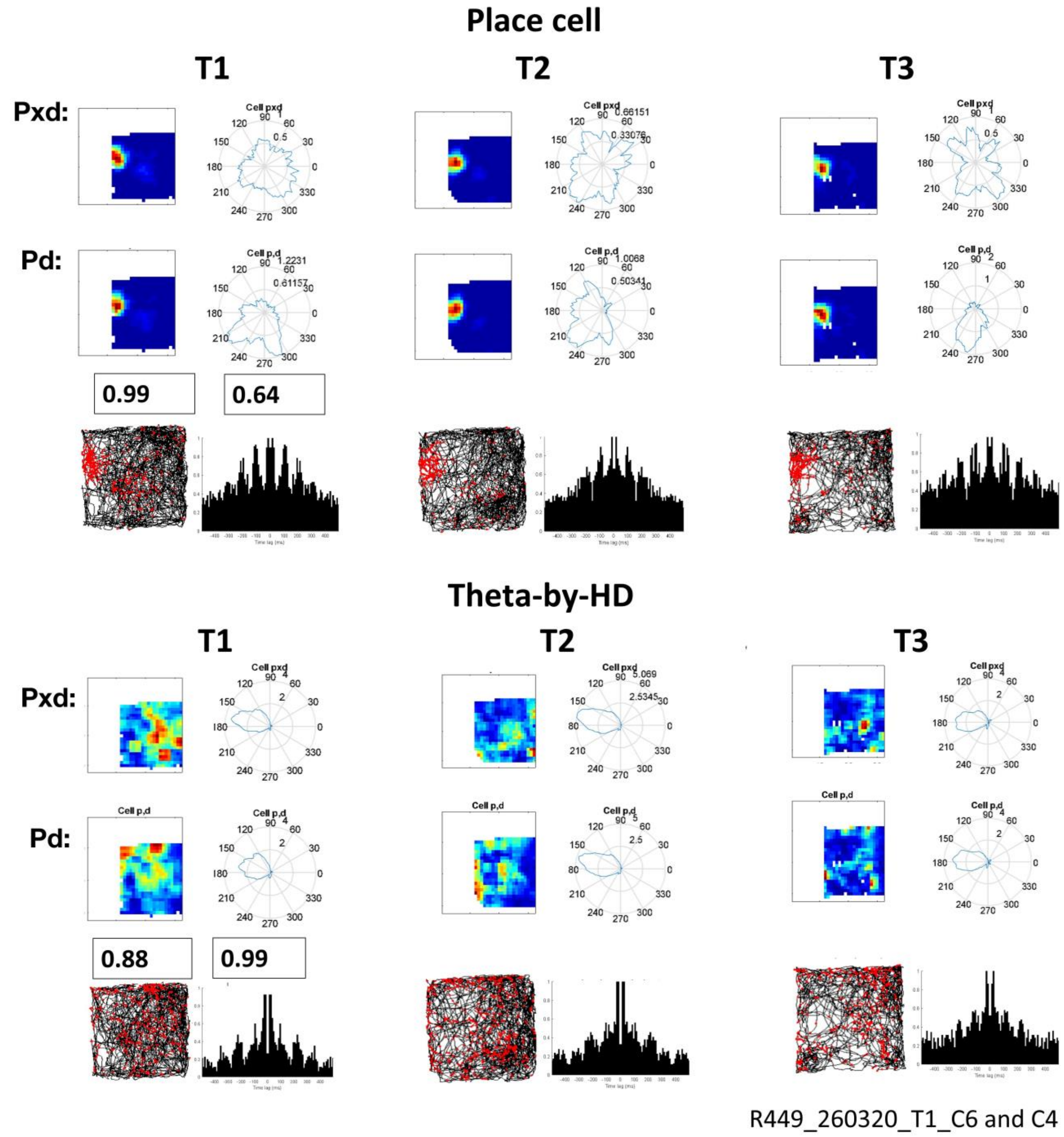


**Figure S8: Effects of applying pxd corrections to the firing plots of a place cell (top) and a Theta-by-HD cell (bottom), A.** Cells were co-recorded on the same tetrode across three trials (each column is a trial, trial name is at the top: T1, T2, T3). For non-conjunctive Theta-by-HD cells, there was a good match between pd and pxd polar plots but not ratemaps. Original firing rate plots are denoted as "pd", corrected plots are denoted as "pxd”. Correlation values are reported for the first trial as a measure of similarity. Spikemap and autocorrelogram implemented between ±500ms are displayed at the bottom for each trial.

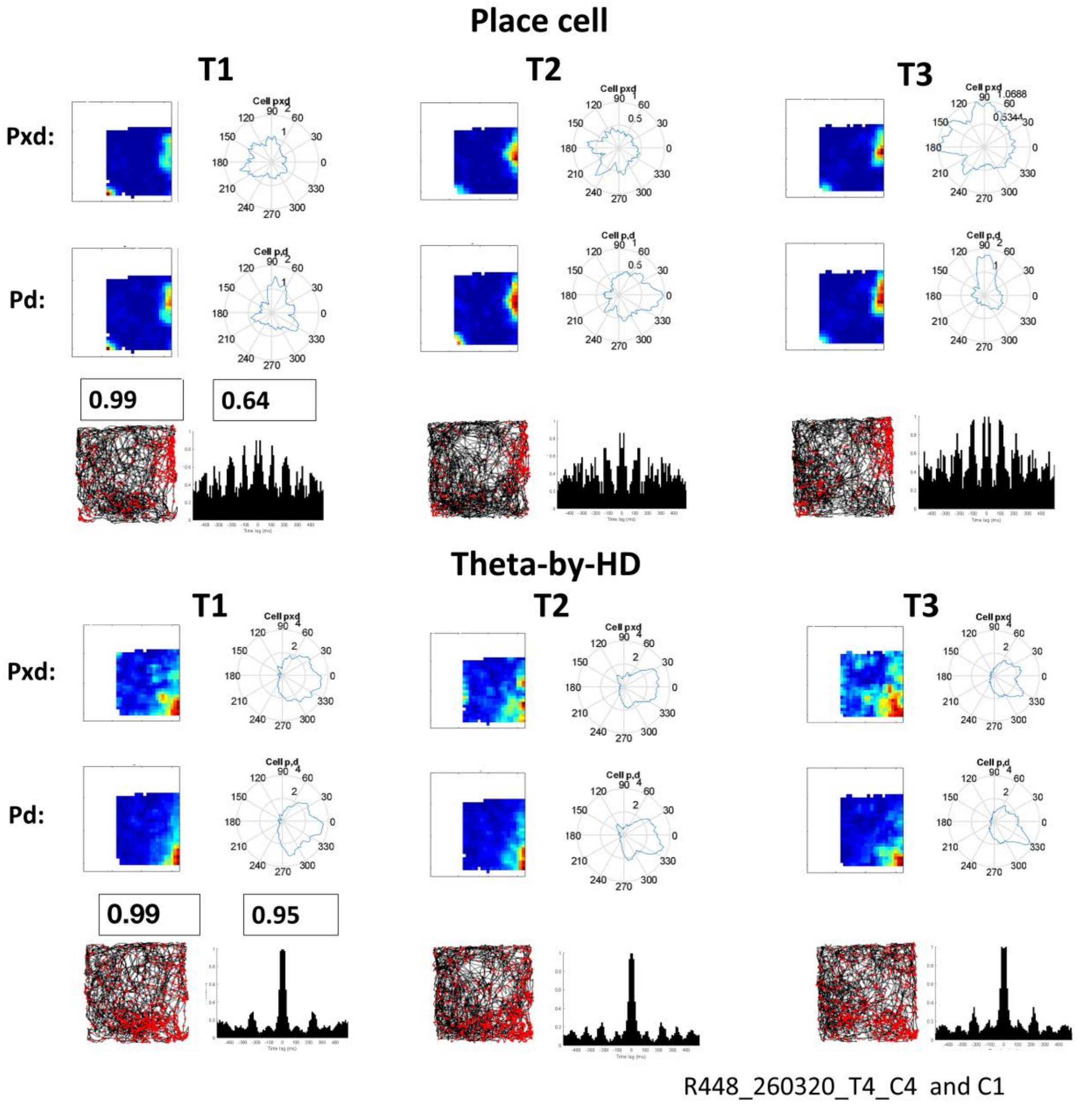


**Figure S9: Effects of applying pxd corrections to the firing plots of a place cell (top) and a Theta-by-HD cell (bottom), B. Cells displayed as per Fig. S7** For place cells, there was a good match between pd and pxd ratemaps but not polar plots. For conjunctive Theta-by-HD cells, there was a good match between pd and pxd polar plots as well as ratemaps.

**Angular head velocity (AHV) had an effect on direction, but not tuning curve shape. Related to Fig. 11**

Tuning curve shape may be influenced by the amount of time by which cells anticipate the animal’s heading direction. Although Theta-by-HD cells have broader directional tuning (see Results), they were less anticipatory and overall showed smaller CW-CCW angular separation compared to non-theta HD cells. Moreover, both cell types retained the same width and height when decomposed into CW and CCW. Hence, broader tuning for Theta-by-HD cells was not a consequence of larger CCW-CW angular separation. Two-way ANOVAs showed that cell type ($F(1,1) = 431.69$, $p < 0.0001$), but not turning direction ($F(1,1) = 0.35$, $p = 0.71$) nor their interaction ($F(1,2) = 0.15$, $p = 0.89$) had an effect on tuning curve width. Similarly, cell type (two-way ANOVA, $F(1,1) = 177.18$, $p < 0.0001$), but not turning direction

(F(1,2) = 0.25, $p$ = 0.78) nor their interaction (F(1,2) = 0.08, $p$ = 0.93) had an effect on peak rate. Values are reported in Table S5. There was a weak negative correlation between a cell's tuning width and its ATI (Spearman's r = -0.16, $p$ = 0.01), and no correlation between a cell's peak firing rate and its ATI (r = 0.12, all $p$ = 0.05).

**Place cell fields do not degrade in darkness. Related to Results: Spatial firing**

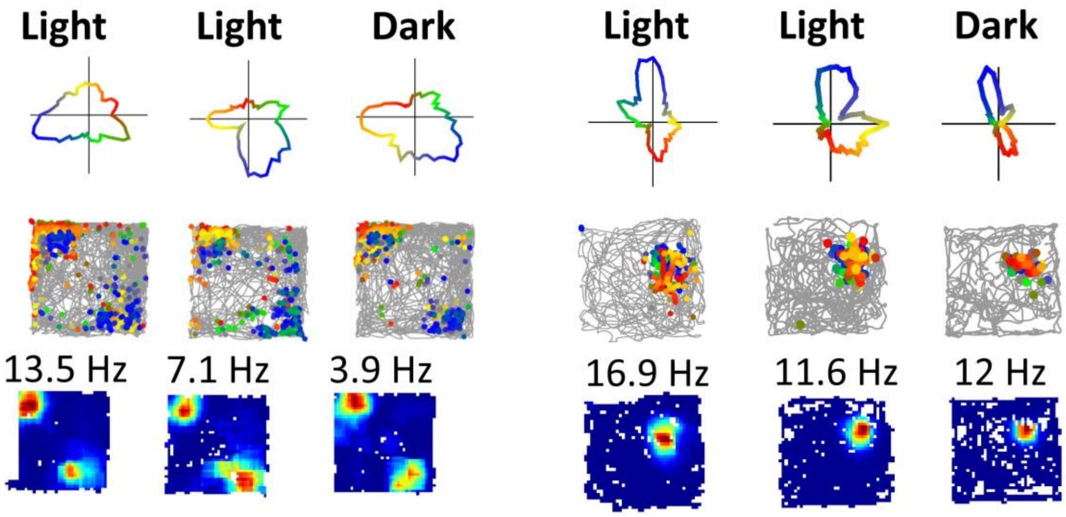


**Figure S10: Spatial firing is similar across Light-Light (Condition 1) and Light-Dark (Condition 2) trials.** Example of two place cells recorded across trials as per Fig. 7C (see main text Results). Absence of vision did not affect place cell representations: ratemap cross-correlations were 0.9 ± 0.01 in Light-Light trials and 0.87 ± 0.03 in Light-Dark trials. No difference between the two Conditions (two-sample KS test; ks-stat = 0.14, $p$ = 0.74).
